# High-Grade Serous Ovarian Tumor Cells Modulate NK Cell Function to Create an Immune-Tolerant Microenvironment

**DOI:** 10.1101/2020.11.20.391706

**Authors:** Veronica D. Gonzalez, Ying-Wen Huang, Shih-Yu Chen, Antonio Delgado-Gonzalez, Kenyi Donoso, Andrew Gentles, Karen Sachs, Ermelinda Porpiglia, Wendy J. Fantl

**Author notes:** 10X Genomics, Pleasanton, CA 94588. Institute of Biomedical Sciences, Academia Sinica, Taipei, 11529, Taiwan (Shih-Yu Chen).

## Abstract

Tubo-ovarian high-grade serous cancer (HGSC) is unresponsive to immune checkpoint blockade despite significant frequencies of exhausted T cells. Here we applied mass cytometry to uncover decidual-like (dl)-NK cell subpopulations (CD56+CD9+CXCR3+KIR+CD3-CD16-) in chemo-naïve HGSC tumors that correlated with both tumor and transitioning epithelial-mesenchymal cell abundance. We showed different combinatorial expression patterns of ligands for activating and inhibitory NK receptors within the three HGSC tumor cell compartments; epithelial (E), transitioning epithelial-mesenchymal (EV) and mesenchymal (vimentin-expressing cells, V) with a more inhibitory ligand phenotype in V cells. When co-cultured with HGSC cell lines the NK-92 cell line acquired CD9 from tumor cells by trogocytosis with a resultant reduction in both anti-tumor cytokine production and cytotoxicity. Critically, a CD9 blocking antibody restored the killing activity of CD9+-NK-92 cells. These findings identify previously unrecognized mechanisms of immune suppression in HGSC. Furthermore, since CD9 is widely expressed in HGSC tumors it represents an important new therapeutic target with immediate relevance for NK immunotherapy.

## Introduction

Tubo-ovarian high-grade serous cancer (HGSC) is the most lethal gynecologic malignancy mainly as the consequence of its advanced-stage diagnosis by which time it has metastasized to multiple sights making curative treatment challenging. Standard-of-care is surgical debulking and platinum-based chemotherapy with a 70 to 80% likelihood of recurrence within 5 years (Bast et al., 2019; Bowtell et al., 2015; Matulonis et al., 2016; Singh et al., 2017). Recently, the introduction of two new treatment modalities into the clinic has brought renewed hope to women with HGSC. One exploits the paradigm of synthetic lethality through the administration of small molecule poly (ADP-ribose) polymerase inhibitors (PARPi). These have been clinically approved for HGSCs harboring loss of function in BRCA1 or BRCA2 genes (Ashworth and Lord, 2018; Lord and Ashworth, 2017). Immunotherapy, the second recently developed treatment modality, is aimed at restoring the ability of the patient’s immune system to eradicate a tumor (Rodriguez and Galpin, 2018) and is an approach mostly focused on reactivation of T lymphocytes. Although HGSC tumors show high frequencies of functionally exhausted T cells, with high levels of immune checkpoint proteins, such as PD-1, CTLA-4, LAG-3 and PD-L1, responses to immunotherapy for HGSC have been disappointing (Kandalaft et al., 2019; Lee and Matulonis, 2019; Rodriguez and Galpin, 2018). Therefore, a deeper understanding of the cell types within the HGSC immune microenvironment could assist in identifying predictive mechanistic biomarkers to select patients likely to gain the most benefit from immunotherapy.

Natural killer (NK) cells are innate lymphocytes with potent cytotoxic activity against tumors and virally infected cells. NK cell function results from the coordinated integration of intracellular signaling mediated by the combinatorial expression of multiple activating and inhibitory germline encoded receptors (Morvan and Lanier, 2016; Orr and Lanier, 2010; Vivier et al., 2011; Wilk and Blish, 2018). NK cells produce an array of cytokines that regulate immune responses, and they are mechanistically distinct from T lymphocytes in that their cytotoxic activity occurs in an antigen-independent manner and without the need for prior sensitization. In tumors these dual functions endow NK cells with roles in both immune surveillance to eradicate tumor cells and conversely with the creation of an immune tolerant microenvironment facilitating tumor progression (Chiossone et al., 2018; Morvan and Lanier, 2016).

In an earlier study of newly diagnosed late stage HGSC tumors by single cell mass cytometry (CyTOF) we demonstrated the presence of three tumor cell compartments comprising epithelial cells (E) expressing E-cadherin, mesenchymal cells (V) expressing vimentin and notably EV cells a phenotype transitioning between the epithelial and mesenchymal compartments (Gonzalez et al., 2018). We now report on the CyTOF data acquired from the T and NK immune cell infiltrate from these tumors.

NK cells are now at the center of a variety of immunotherapeutic approaches to exploit their tumor cell killing activity (Daher and Rezvani, 2018; Li et al., 2018; Lorenzo-Herrero et al., 2018; Rezvani, 2019; Rezvani et al., 2017; Uppendahl et al., 2017). The single cell data from this study identifies critical and unappreciated mechanisms by which HGSC cells can subvert the killing activity of NK cells. Moreover, this needs urgent consideration when optimizing NK cell-based immunotherapy.

## Results

### CyTOF analysis identifies T and NK cell clusters from HGSC tumor infiltrates

In our prior CyTOF study of seventeen newly diagnosed HGSCs (Tables S1 and S2), computational analysis focused on the tumor cells (Gonzalez et al., 2018). We identified 56 tumor cell phenotypes through an unsupervised analysis of the CyTOF datasets with the X-shift clustering algorithm (Samusik et al., 2016). Expression patterns of E-cadherin and vimentin, critical proteins in epithelial tumors, enabled the identification of three tumor cell compartments; i) epithelial tumor cells that exclusively expressed E-cadherin (E compartment) ii) mesenchymal tumor cells that exclusively expressed vimentin (V compartment) and iii) seven transitional epithelial-mesenchymal tumor cell clusters that co-expressed E-cadherin and vimentin (EV compartment) (Gonzalez et al., 2018; Zhang and Weinberg, 2018). Critically, a subset of V cells correlated with poor prognosis (Gonzalez et al., 2018).

Here we report our further analysis of these HGSC tumors with a CyTOF antibody panel designed to characterize T and NK cell subtypes (Table S3, STAR Methods). All steps for quality control and CyTOF processing of barcoded samples were as previously described (Gonzalez et al., 2018)) (STAR Methods). For each tumor, the immune cell infiltrate was gated out as CD45+CD66-from the viable immune cell population (STAR Methods). The resultant T and NK immune single cell data sets were then combined and subjected to unsupervised analysis with X-shift clustering (Samusik et al., 2016). Using 25 surface markers (Table S3) delineating T and NK cell subpopulations, 52 X-shift cell clusters (calculated for an optimal “k” of 30 nearest neighbors) were generated for subsequent analysis (Samusik et al., 2016).

### Correlation analysis between tumor and immune cell clusters from HGSC tumors

In order to understand the interactions between tumor and immune cells we applied a network approach to compute correlations between the frequencies of these cell phenotypes (Gonzalez et al., 2018; Hotson et al., 2016; Ideker and Krogan, 2012; Spitzer et al., 2017). The resultant pair-wise Spearman correlation coefficients (r_s_) were displayed on a hierarchically clustered heat-map (Figure 1A). For each HGSC tumor, pairwise correlations included the following parameters i) Cell frequencies for all 52 T and NK cell clusters, ii) Cell frequencies for all 56 tumor cell clusters (Gonzalez et al., 2018) iii) Total tumor cell frequency iv) Total E cell frequency v) Total V cell frequency vi) Total EV cell frequency and vii) other features described previously (Gonzalez et al., 2018).

**Figure 1:**
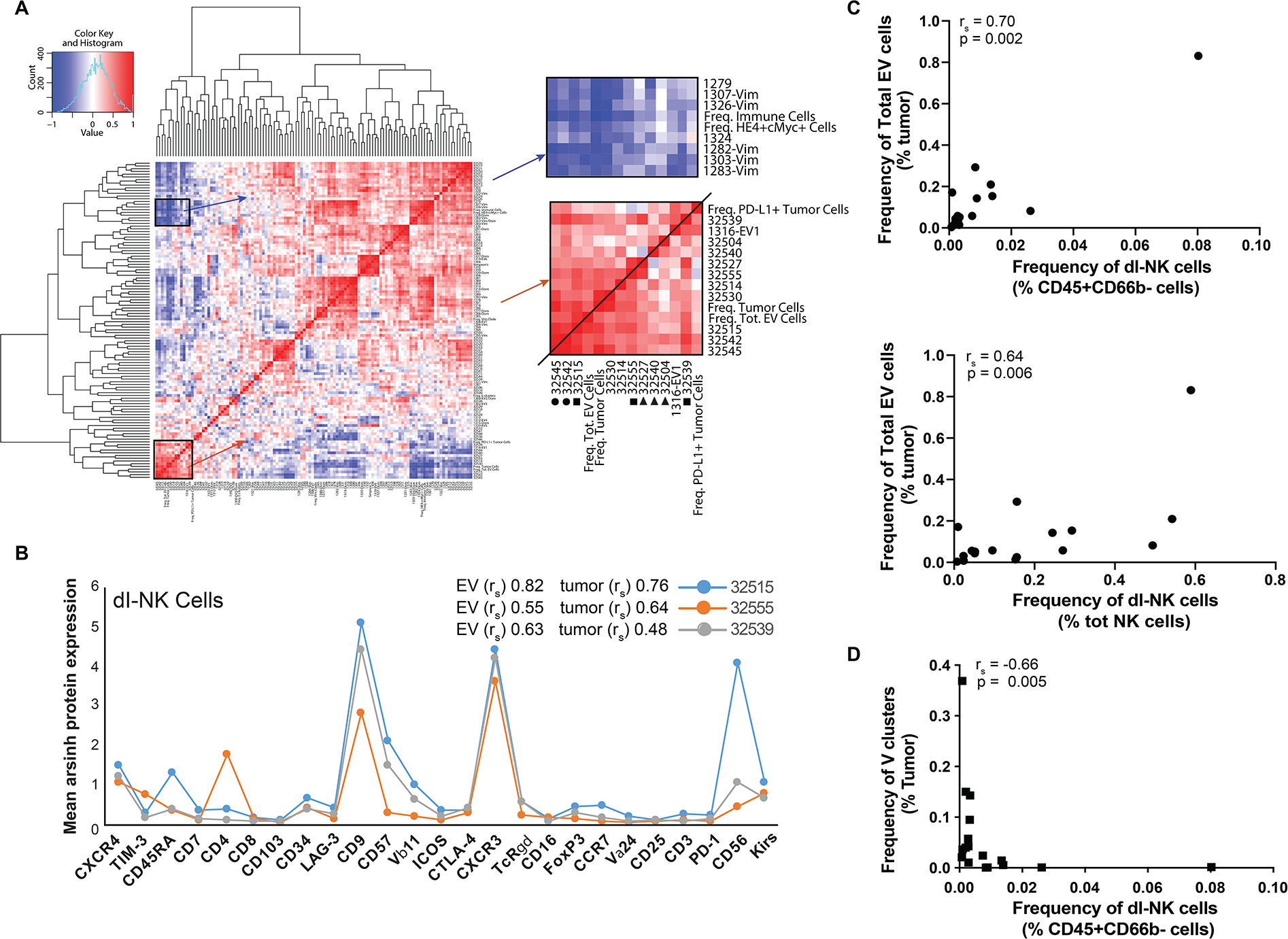
HGSC tumor and EV cell frequencies correlate with a dl-NK cell phenotype. **(A)** Hierarchically organized heat-map showing pairwise Spearman correlations between total tumor and total EV cell abundance with specific immune cell clusters. Enlarged portions of the heat map (right-hand side) depict positive (red) and negative (blue) r_s_ correlations, respectively with dl-NK cell clusters. Positive correlations with total abundance of tumor and EV cells respectively for dl-NK cell clusters (■) and T cell clusters (●) dl-NK cell clusters present in all tumors but showing no correlations (▲). **(B)** Protein expression patterns and correlations of three dl-NK cell clusters. **(C)** Correlation plots of EV cell abundance and percentage dl-NK cells manually gated from the immune cell infiltrate (upper plot) or total NK cell population (lower plot). **(D)** Plot of percentage of manually gated dl-NK cells in immune cell infiltrate with a subgroup of vimentin clusters. See also Figures S1 - 3 and Tables S1 – S3.

Analysis revealed five immune cell clusters that correlated with both total tumor and EV cell abundance (Spearman’s correlation coefficient, r_s_ > 0.5) (Figure 1A). Three of these immune cell clusters (32515, 32539 and 32555) were negative for CD3 and CD16 expression but positive for expression of CD56, CD9, CXCR3 and killer immunoglobulin-like receptors (KIRs), a phenotype that resembles decidual NK cells (Koopman et al., 2003) (Figure 1B). Notably, among the many NK cell phenotypes reported, CD9 expression is exclusive to decidual NK (d-NK) cell subsets (Horowitz et al., 2013; Koopman et al., 2003; Wilk and Blish, 2018). The combined cell frequency of these three decidual-like NK (dl-NK) clusters ranged from 1.3% - 28% across the 17 tumors. Furthermore, correlations determined with manually gated dl-NK cells and EV tumor cells were in agreement with the unsupervised X-shift analysis (Figure 1C and Figure S1).

The remaining two immune cell clusters (32542 and 32545) had a T-cell phenotype (CD3+ with mutually exclusive expression of CD4 and CD8) but also shared phenotypic properties with decidual NK cells. They had high levels of CD56, CXCR3 and CD9 and low levels of the invariant T cell receptor Vα24-Vβ11 suggesting that these cells could have NKT-like functions (Bernstein et al., 2006; Kim et al., 2002; Koreck et al., 2002; Reyes et al., 2018) (Figure S2A). They were present in the HGSC immune cell infiltrate with a frequency of 0.1 – 3.9%.

An additional three dl-NK cell clusters (32527, 32504 and 32540) were found that did not correlate with either tumor or EV cell abundance (Figure 1A and Figure S2B). Cluster 32527 correlated with dl-NK cell clusters 32555 and 32539 (Figure 1B). Clusters 32504 and 32540 did not correlate with any tumor features but were present in all tumors with a combined frequency of 14% - 86% of the immune cell infiltrate. To visualize the relationships between all T and NK immune cell clusters, a minimum spanning tree was generated (Bendall et al., 2011; Gonzalez et al., 2018). This revealed that clusters 32504 and 32540 were phenotypically similar to both dl-NK and T cell clusters (Samusik et al., 2016) (Figure S3).

Of the eight immune cell clusters described above, five correlated negatively (r_s_ > −0.6) with several vimentin clusters (Figure 1D). These data are consistent with published reports describing an inverse relationship between metastases and NK cells within immune infiltrates (Lopez-Soto et al., 2017; Lorenzo-Herrero et al., 2018).

Decidual NK cells play a critical role in the first trimester of pregnancy by conferring immune tolerance toward the hemi-allogeneic fetus and facilitating placental growth (Cooper et al., 2001; Hanna et al., 2006; Hanna and Mandelboim, 2007; Jabrane-Ferrat, 2019; Koopman et al., 2003). The identification of dl-NK cells in HGSC led us to hypothesize that the same features of dNK-mediated immune tolerance could be subverted for HGSC tumor maintenance and progression.

### NK receptor ligand expression across newly diagnosed HGSC tumors

Having previously identified the E, EV and V intra-tumor cell compartments, each representing different stages of disease progression we wished to determine how these compartments modulated NK cell function toward an immune-tolerant state (Gonzalez et al., 2018). We analyzed CyTOF data from 935,563 single intact cells, prepared from 12 newly diagnosed late stage HGSC tumors, and determined frequencies of cells expressing NK receptor ligands and two ADAMs (a disintegrin and metalloproteinase) across the three tumor compartments (Table S1, STAR Methods). The modified HGSC CyTOF panel included antibodies against the following twelve NK receptor ligands and two ADAM proteases (Table S4) (Gonzalez et al., 2018; Zhang and Weinberg, 2018): i) ULBP1, ULBP2/5/6, ULPBP3, ULPBP4 and MICA/B that bind to the NKG2D activating NK receptor (Dhar and Wu, 2018; Raulet et al., 2013), ii) ADAM10 and ADAM17 proteases involved in NK ligand and NK receptor shedding (Boutet et al., 2009; Dhar and Wu, 2018; Ferrari de Andrade et al., 2018), iii) nectin-like ligands, CD111, CD112, CD113, CD155 and nectin-4. These bind the activating NK receptor, CD226 (also known as DNAM1) and the inhibitory NK receptors, T cell immune-receptor with immunoglobulin and ITIM domains (TIGIT) and CD96 (also known as TACTILE) (Fabre-Lafay et al., 2007; Martinet and Smyth, 2015; Reches et al., 2020; Sanchez-Correa et al., 2019), iv) Human leukocyte antigen (HLA) class 1 molecules, A, B and C that bind to inhibitory killer immunoglobulin-like receptors (KIR) (Morvan and Lanier, 2016; Wroblewski et al., 2019) and, v) HLA-E, a non-classical HLA class 1 molecule, that binds to the NK inhibitory receptor heterodimer CD94/NKG2A with greater affinity than to the activating CD94/NKG2C heterodimer (Kamiya et al., 2019).

### NK receptor ligand expression levels across tumor cell compartments

In order to visualize expression levels of the NK receptor ligands on tumor cells, CyTOF datasets for each of the twelve HGSC samples were manually gated to exclude immune, angiogenic and stromal cells (CD45-, CD31-, FAP-) (Gonzalez et al., 2018). The resultant single cell data files were combined and tumor cells were clustered as before using the X-shift algorithm (Table S4) (Gonzalez et al., 2018; Samusik et al., 2016).

Spatial relationships between tumor cells expressing NK receptor ligands within the 56 X-shift tumor cell clusters were visualized by force directed layouts (FDLs). 10,000 single cells were computationally sampled from each X-shift cluster and each cell was connected on a 10-nearest-neighbor graph. This graph was subjected to a force-directed layout that placed groups of phenotypically related cells adjacent to one another (Samusik et al., 2016) (Figure 2A – C, STAR Methods). The resultant FDLs were composites of tumor cells from all twelve samples and corroborated the presence of the E, V and EV compartments (the last comprised of seven clusters) as previously reported (Gonzalez et al., 2018) (Figure 2A, upper two panels with V and EV cells encircled). In general, the FDLs revealed that receptor ligands, for both activating and inhibitory NK receptors, were expressed at variable levels in all three compartments within pockets of tumor cells, rather than evenly interspersed throughout the tumor cells (Figure 2A - C).

**Figure 2:**
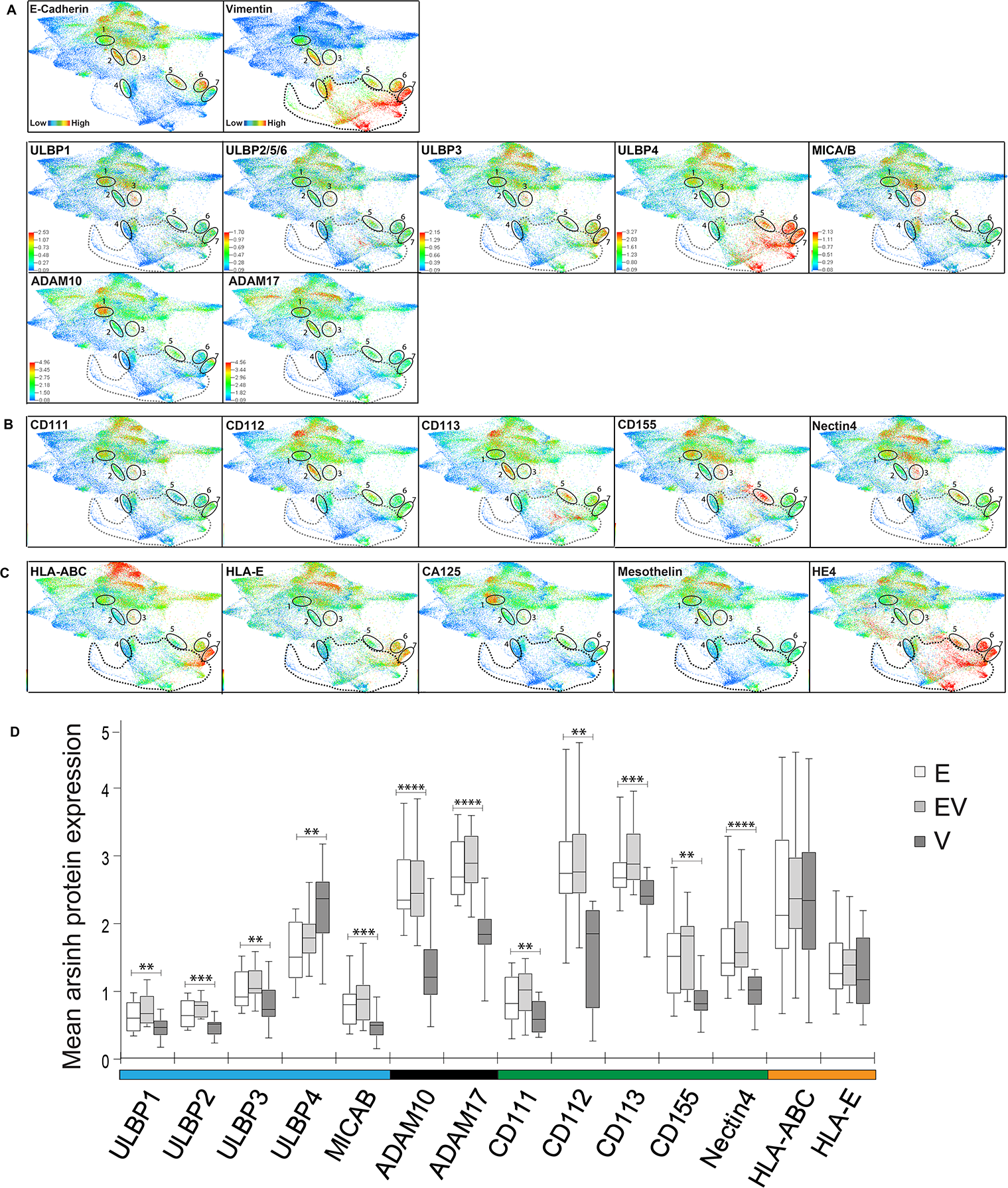
Expression patterns of NK receptor ligands in newly diagnosed HGSC tumors. Single cell force directed layouts (FDLs) are composites of twelve HGSC tumors. Single cell sub-sampling with generation of FDLs was repeated three times with comparable results. Panels for **(A)** E-cadherin and vimentin (top). EV clusters co-expressing E-cadherin and vimentin are encircled and numbered 1 to 7 (Gonzalez et al., 2018), NKG2D activating receptor ligands (second row) and ADAM proteases (third row). **(B)** Nectin-family ligands **(C)** HLA-ABC and HLA-E inhibitory ligands and tumor associated antigens; CA125, mesothelin and HE4. **(D)** Box and whisker plots. Expression levels for 12 NK receptor ligands and two ADAM proteases across E, EV and V compartments of 12 HGSC tumors. Medians and interquartile ranges are shown. p-values: ** ≤ 0.01, *** ≤ 0.001, **** ≤ 0.0001. See also Table S4.

Previous studies have demonstrated that upregulation of ligands for the NKG2D activating NK receptor is a major mechanism by which NK cells are able to detect and eradicate tumor cells (Dhar and Wu, 2018; Raulet et al., 2013). Our data revealed that, apart from ULPBP4, the highest levels of activating NKG2D receptor ligands were found in the E and EV HGSC tumor compartments with minimal levels in the V compartment (Figure 2A). This is consistent with an immune-surveillance role for NK cells within the E and EV tumor cell compartments. The high levels of ULBP4 (which also binds to NKG2D) throughout the V cell compartment are an anomaly but consistent with a recent study describing ULBP4 as a functional outlier within the ULBP family (Zoller et al., 2018).

For the proteases ADAM10 and ADAM17, discrete pockets of cells with high expression levels were observed in the E compartment and in the EV1 transitional tumor cell subset. Pockets of ADAM-expressing tumor cells co-localized with NKG2D ligands suggesting an attempt by tumor cells to nullify NK cell killing activity by promoting NK ligand shedding (Dhar and Wu, 2018; Ferrari de Andrade et al., 2018; Raulet et al., 2013).

The nectin family of ligands also exhibited variable expression patterns with pockets of E tumor cells, co-expressing high levels of CD112, CD113 and CD155. EV2, EV3 and EV5 co-expressed different combinations of CD112, CD113, CD155 and nectin 4 (Figure 2B).

HLA-A, B, C and E primarily engage NK cell inhibitory receptors to provide self-tolerance for healthy cells (Morvan and Lanier, 2016). These ligands were also co-expressed in pockets of cells at varying levels in all three tumor compartments thus suppressing tumor cell destruction by NK cells (Figure 2C). By contrast, MHC class I molecules expressed by tumor cells, targets them for destruction by cytotoxic CD8 T lymphocytes (CTLs) and occurs in an antigen-dependent mechanism whereby MHC class I molecules present peptide fragments from tumor associated antigens (TAAs) to CTLs (Brennick et al., 2017; Vyas et al., 2008). Well-established TAAs in HGSC are MUC16 and mesothelin (Bast et al., 2019). For the most part expression of these TAAs and HLAs were mutually exclusive indicating minimal tumor cell destruction by CTLs (Figure 2C). These data suggest that HGSC cells may have evolved dual mechanisms of escape from the killing activity of both NK and CD8 T cells.

### Different expression levels of NK receptor ligands and ADAMs across E, EV and V tumor compartments

Box and whisker plots revealed that median expression levels for the NK receptor ligands and ADAMs did not differ significantly between the E and EV compartments (Figure 2D). However, with the exception of ULBP4, within the V compartment they were all statistically lower (Table S5) enabling V cells to more easily escape immune surveillance; a result consistent with previously reported experimental models (Lopez-Soto et al., 2017; Lorenzo-Herrero et al., 2018). There were no statistical differences between tumor compartments for the expression of HLA-ABC and E.

### Quantifying the combinatorial diversity of NK receptor ligand expression

The combinatorial expression patterns of activating and inhibitory receptors endow NK cells with a high degree of phenotypic and functional diversity (Horowitz et al., 2013; Wilk and Blish, 2018). In their turn, a correspondingly complex repertoire of activating and/or inhibitory ligands present on tumor cells can affect NK receptor activity (Morvan and Lanier, 2016). In order to determine differences in NK receptor ligand expression combinations within the immune microenvironment of the E, EV and V tumor compartments, we applied Boolean analysis to measure the frequency of cells with distinct combinatorial expression patterns for the twelve NK receptor ligands and the two ADAM proteases. We assessed 2^14^ (16,348) NK receptor ligand combinations and compared the frequency of tumor cells harboring specific NK receptor ligand combinations (in the E, EV and V tumor compartments) (Figure 3A, STAR Methods). Using a threshold cell frequency of >1% for cells in any compartment in any sample expressing an NK receptor ligand/ADAM protease combination there were 163 NK receptor ligand combinations expressed by tumor cells in the 12 HGSCs (Figure 3A, ligand combinations in rows).

**Figure 3:**
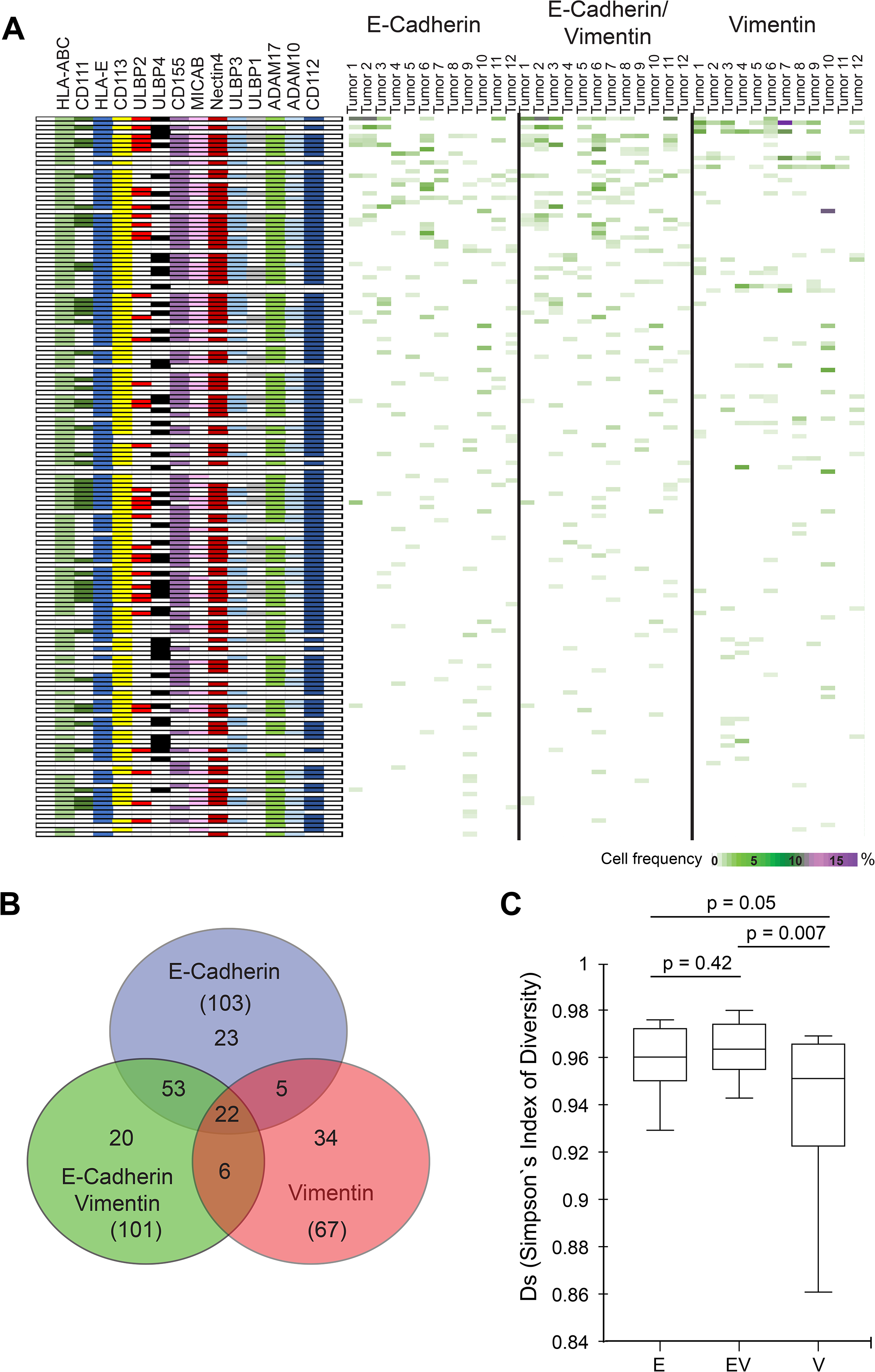
Combinatorial diversity for NK receptor ligands within E, EV and V HGSC tumor compartments. **(A)** Boolean logic computation (left panel). Combinatorial diversity of 12 NK receptor ligands and ADAM 10 and 17 proteases expressed by tumor cells. Each NK receptor ligand combination is a row (left hand side). The heat map (right panels) shows the frequency of tumor cells that express each ligand combination within the E, EV, and V compartments for each tumor sample (columns). Rows were ranked based on the highest (top) to lowest total cell frequency (bottom). **(B)** Venn diagram showing distinct and overlapping NK receptor ligand combinations across E, EV and V compartments. **(C)** The Simpson’s inverse index of diversity was significantly greater for the E versus V (p = 0.05) and EV versus V (p = 0.007) compartments across all HGSC tumors. Medians and inter quartile ranges are shown.

Boolean analysis demonstrated that cells in the E and EV tumor compartments had greater combinatorial diversity for NK receptor ligands/ADAMs (103 and 101 phenotypes respectively) than the V tumor compartment (67 phenotypes) (Figure 3B). There were more shared phenotypes between the E and EV compartments (53) than between the E and V (five) and EV and V (six) compartments (Figure 3B). These data suggest differential regulation of the immune microenvironment orchestrated by the tumor cells in each of the three compartments.

To quantify further the NK receptor ligand combinatorial diversity in the different tumor cell compartments we calculated the Simpson’s index of diversity. (Figure 3C). This index is often used in ecology to quantify the biodiversity within a natural habitat and was recently applied to NK and ovarian tumor CyTOF datasets (Gonzalez et al., 2018; Horowitz et al., 2013). When applied here, the Simpson’s index of diversity was significantly higher in the E and EV compartments compared to the V tumor compartment consistent with the greater number of NK receptor ligand combinations in the E and EV compartments compared to those in the V compartment (Figure 3B).

### NK receptor ligand expression across HGSC cell lines

In their genetic analysis of ovarian cancer cell lines Domcke et al. presented a group of ovarian cell lines ranked by the concordance of their genetics to resected HGSC tumors, with the goal of providing more reliable in vitro models of HGSC (Domcke et al., 2013). In order to determine how phenotypes of these cell lines compared with HGSC tumor cells, we analyzed thirteen of the highest ranked HGSC cell lines with our CyTOF tumor antibody panel modified with antibodies against NK receptor ligands and ADAMs (Table S4). Data analysis performed with X-shift clustering and FDL visualization, revealed HGSC cell lines that phenocopied E, EV and V tumor compartments, based on their expression of E-cadherin and vimentin. (Figure S4A). Examination of the NK receptor ligand/ADAM expression levels across the E, V and EV cell lines revealed expression patterns that, although not identical, were comparable to their E, V and EV counterparts in HGSCs (Figure S4). For example, the NKG2D activating ligands were expressed primarily in E and EV cell lines and at background levels in V cell lines (Figure S4B and E). Based on these observations we selected three HGSC cell lines for in vitro studies, OVCAR4, Kuramochi and TYK-nu, representing E, EV and V HGSC tumor cells, respectively.

### Carboplatin changes NK receptor ligand expression levels

It is well established that activation of the DNA damage response with genotoxic agents increases the expression of ligands for NKG2D and DNAM1 thereby making a “stressed” cell more susceptible to NK cell killing (Cerboni et al., 2014; Gasser et al., 2005). We therefore exposed the three HGSC cell lines above to carboplatin, a genotoxic agent that is part of the standard-of-care regimen for women with HGSC (Bast et al., 2019; Bowtell et al., 2015; Matulonis et al., 2016). After one week, the HGSC cell lines were processed for CyTOF using the tumor/NK receptor ligand antibody panel (Table S4).

The DNA damage response after carboplatin treatment was confirmed by a recognized increase in pH2AX (Krenning et al., 2019) (Figure 4). However, of the NK activating receptor ligands measured, only ULBP2 increased in OVCAR4 cells in response to carboplatin (24% to 50%). Evaluation of ligands for inhibitory NK receptors revealed significant increases of cells expressing HLA-E in all three cell lines and a significant increase in Kuramochi cells expressing HLA-ABC.

**Figure 4.**
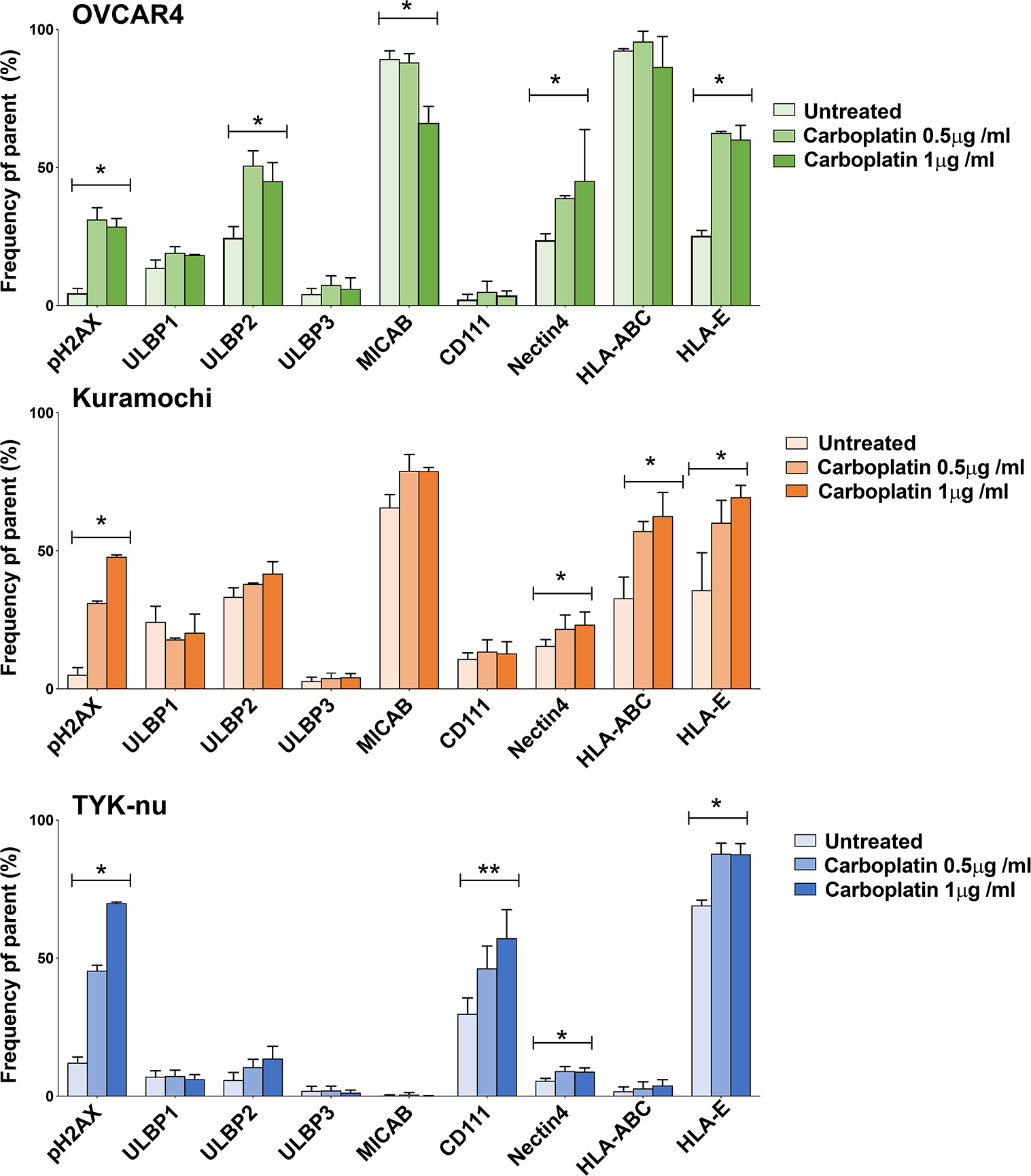
Responses to carboplatin across E, EV and V HGSC cell lines. OVCAR4 (E), Kuramochi (EV) and TYK-nu (V) cell lines exposed to vehicle or carboplatin at 0.5 or 1μg/mL for 1 week were processed for CyTOF with the tumor NK receptor ligand/ADAM antibody panel (Table S3). Parent populations (viable single cells negative for cisplatin and cPARP). The plots show frequencies of HGSC cells expressing activating and inhibitor NK receptor ligands (X-axis) gated out of the parent population (Y-axis). Plots show the mean of triplicates with standard deviations. *p ≤ 0.05, **p ≤ 0.005, for overall ANOVA.

Carboplatin also increased the frequency of HGSC cells expressing nectin 4, particularly OVCAR4. Prolonged exposure of cells to carboplatin (at two different doses) increased the frequency of nectin-4-positive OVCAR4 cells from 24% (baseline) to 39%, and 45%. Similarly, for Kuramochi from 15% to 22%, and 23% with modest changes in TYK-nu cells from 6%, to 9% and no increase thereafter (Figure 4). In trophoblasts overexpression of nectin 4 increases susceptibility to NK cell-mediated cytotoxicity (Ito et al., 2018). Thus, by analogy after carboplatin exposure OVCAR4 showed the greatest susceptibility to NK cell-mediated cytotoxicity with apparently decreasing susceptibility for Kuramochi and TYK -nu cells (Figure 4). In TYK-nu cells, carboplatin mediated a significant increase in CD111, a ligand for the inhibitory CD96 receptor. CD111 also has a role in enhancing signaling through TIGIT (Martinet and Smyth, 2015). These carboplatin-mediated changes toward a more inhibitory NK receptor ligand phenotype reveal an as yet unrecognized mechanism of resistance to this agent.

### HGSC – NK-92 cell line cocultures to model the HGSC immune tolerant microenvironment

The positive association between dl-NK cell sub-populations and overall tumor and EV cell abundance indicate that dl-NK cells are immune tolerant toward the tumor, allowing its continued growth (Figure 1A – C). To determine how the tumor cells might create this immune tolerant environment we set up co-cultures between OVCAR4, Kuramochi and TYK-nu HGSC cell lines with the human NK-92 cell line (Koopman et al., 2003; Suck et al., 2016). After co-culture we performed CyTOF analysis with an NK cell CyTOF antibody panel (Table S6). We chose the NK-92 cell line because of its clinical development for adoptive cellular NK immunotherapy (Daher and Rezvani, 2018; Rezvani, 2019; Rezvani et al., 2017).

### CD9 expression in NK-92 cells dramatically increases after coculture with HGSC cell lines

In monoculture, NK-92 cells showed background CD9 expression. However, after coculture with HGSC cell lines, up to 60% of NK-92 cells expressed CD9 with the greatest induction seen with OVCAR4 cells. (Figure 5A). When the coculture was performed with a membrane barrier (transwell) between the two cell lines, CD9 expression on NK-92 cells was dramatically reduced (< 4%) demonstrating the requirement for physical contact between HGSC tumor cells and NK-92 cells (Figure 5A).

**Figure 5:**
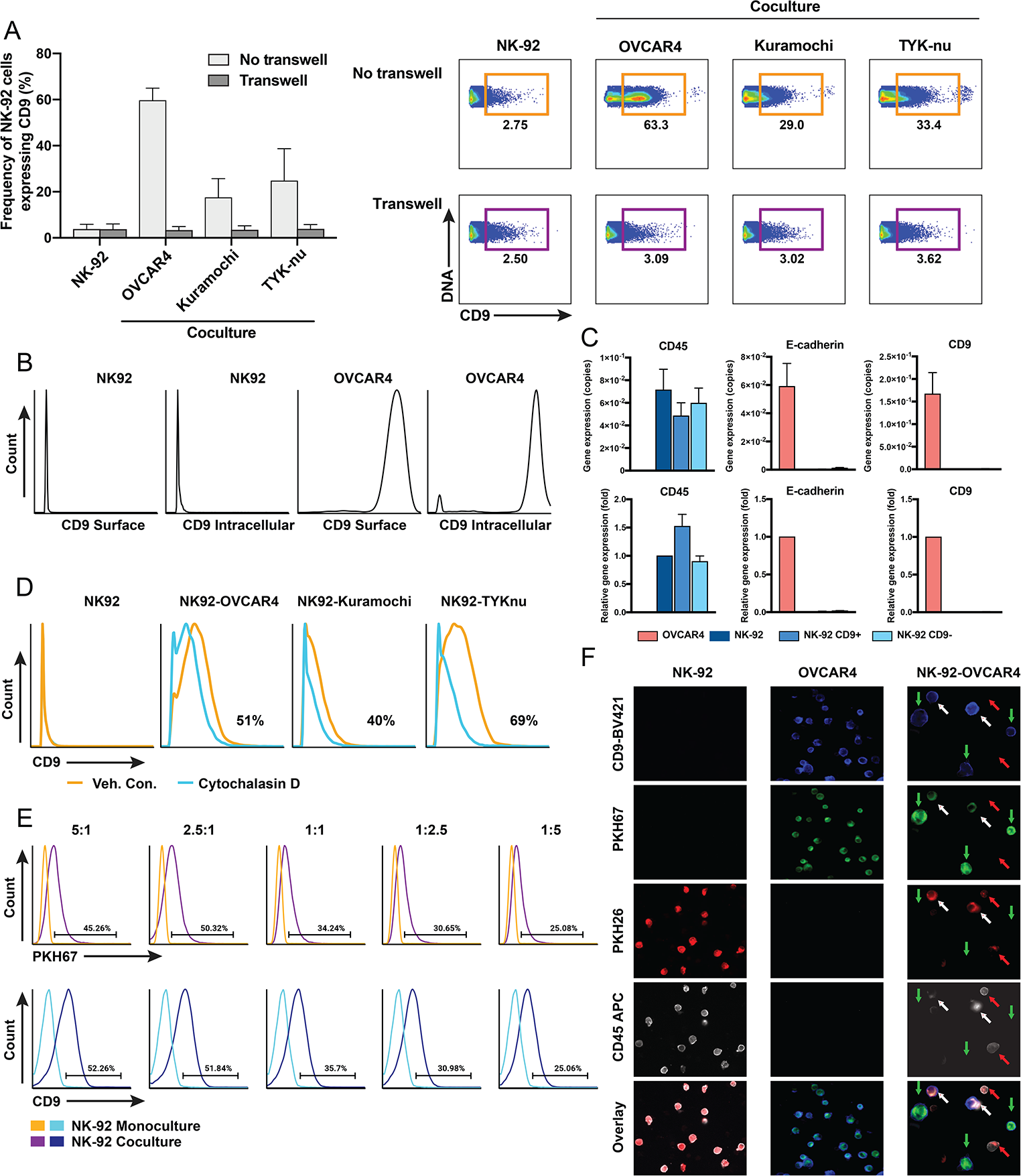
Trogocytosis from HGSC to NK-92 cell lines. HGSC and NK-92 cell lines were cocultured at an effector (NK-92): target (OVCAR4) ratio of 1:1 for 6h unless otherwise indicated (STAR Methods). **(A)** Left panel. Frequency of CD9 expression in NK-92 cells post coculture with and without transwell, respectively. Mean with standard deviations are shown (n=4). Right panel. Exemplary 2D flow plots. Induction of CD9+ NK-92 cells after coculture. **(B)** Lack of extra- and intracellular CD9 protein expression in the NK-92 cell line but high extra- and intracellular CD9 protein levels in the OVCAR4 cell line. **(C)** RT-PCR of FACS purified CD9+ and CD9-NK-92 cells after coculture with OVCAR4 cells. Number of copies (upper plots) or fold gene expression changes (lower plots) after coculture compared to respective monocultures. **(D)** Pre-incubation of NK-92 cells with cytochalasin D (10μM), a trogocytosis inhibitor, before coculture with HGSC cell line results in partial inhibition of trogocytosis. **(E)** Transfer of membrane fragments with CD9 from OVCAR4 cells stained with PKH67 onto NK-92 cells. Cocultures shown at different target : effector ratios. PKH67 frequency changes (upper histograms) and CD9 percentage changes (lower histograms). **(F)** Visualization of trogocytosis by microscopy. OVCAR4 cells and NK-92 cells stained with PKH67 (green) and PKH26 (red) respectively, were cocultured for 3h and stained with antibodies against CD45 and CD9. Images (from Keyence BZ-X800 microscope) for cells grown in monoculture 20X and for coculture 60X. Arrows mark cells NK-92 cells (red), OVCAR4 cells (green) and NK-92 cells after trogocytosis (white). Images were enhanced for brightness and contrast to optimize the printed image. See also Figure S6.

One potential explanation for the appearance of surface CD9 expression on NK-92 cells is that they retain intracellular CD9 pools that during coculture with HGSC cells, are induced to traffic to the cell surface. To address this, we stained NK-92 and OVCAR4 cells grown in monoculture with a CD9 antibody (STAR Methods). Sequential cell staining for CD9 (surface and, following permeabilization intracellular) showed that NK-92 cells were devoid of both surface and intracellular CD9. By contrast, OVCAR4 cells expressed robust levels of CD9 in both cellular locations (Figure 5B and STAR Methods).

To further confirm that CD9 displayed on NK-92 cells during co-culture was not endogenously produced, CD9+ NK-92 cells and their negative counterpart CD9-NK-92 cells were FACS sorted after coculture with the OVCAR4 cell line and analyzed for CD9 transcripts (Figure 5C). Transcripts were not detected in either CD9+ or CD9-FACS sorted NK-92 cells. By contrast, robust levels of CD9 transcripts, consistent with CD9 protein expression, were seen in the OVCAR4 line with which NK-92 were co-cultured. Control transcripts measured were CD45 (positive for NK-92; negative for OVCAR4) and E-cadherin (negative for NK-92; positive for OVCAR4) (Figure 5C).

### CD9 expression across HGSC primary tumors and cell lines

In order to determine how prevalent CD9 expression was in HGSC we screened 17 primary HGSC tumors and 11 HGSC cell lines to determine both the frequency of CD9-expressing cells and the levels of CD9 expression (Tables S1 and S7). For the primary tumor cohort, high frequencies of CD9+ tumor cells were present in all 17 samples (Figure S5). For HGSC cell lines, we screened the top ranking HGSC cell lines as reported by Domcke et al. and observed overall high frequencies of CD9-expression (Domcke et al., 2013) (Figure 6A, B).

**Figure 6:**
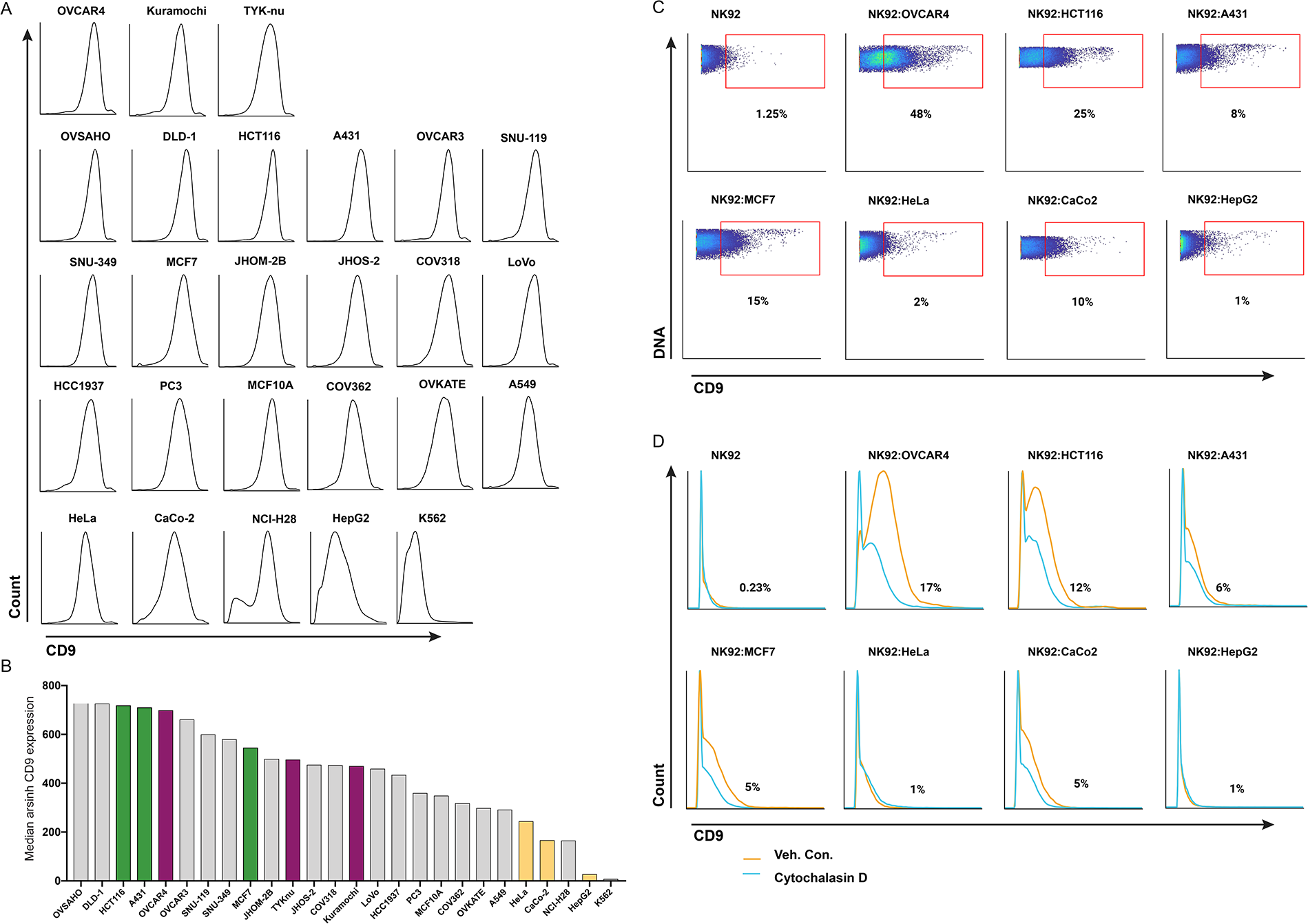
Trogocytosis of CD9 from non-HGSC cells. **(A)** Eleven HGSC and 15 non-HGSC tumor cell lines were screened by CyTOF for CD9 expression. **(B)** Cell lines ranked by level of CD9 expression. Cell lines selected for coculture with NK-92 cells; HGSC (E, EV and V) (magenta), non HGSC cell lines with high levels of CD9 (green) and non-HGSC cell lines with lower levels of CD9 (yellow). **(C)** Representative flow plots showing the frequency of CD9+ NK-92 cells after coculture with non-HGSC cell lines. OVCAR4 included as a control. **(D)** Preincubation of NK-92 cells with cytochalasin D (10μM) and coculture with cell lines as in (C) results in partial inhibition of trogocytosis.

### Trogocytosis as the mechanism by which NK-92 cells acquire CD9

Since there was no evidence of de novo synthesis of CD9 by NK-92 cells we hypothesized that during co-culture CD9 must be transferred to NK-92 cells by a process known as trogocytosis (Dance, 2019; Joly and Hudrisier, 2003). This process takes place within minutes of cell-cell contact and involves the transfer of plasma membrane fragments from one cell to another, including any anchored proteins therein. After coculture with OVCAR4 cells (15, 30, 60, 120 and 360min), CD9 was detected as early as 15min on NK-92 cells with a steady increase up to 360min (Figure S6). These data are consistent with trogocytosis as previously reported in other systems (Reed and Wetzel, 2019).

Many studies have shown that inhibitors of actin polymerization block trogocytosis but that the effect of these inhibitors vary depending on cell type (Aucher et al., 2008; Gary et al., 2012). Our pilot experiment tested a series of trogocytosis inhibitors; concanamycin A, LY294002, EDTA, nocodazole and cytochalasin D. Cytochalasin D was selected as the optimal inhibitor (STAR Methods). Its inclusion in coculture experiments with NK-92 and OVCAR4, Kuramochi or TYK-nu HGSC cell lines resulted in 40 – 69% reduction of CD9+ NK-92 cells (Figure. 5D).

To establish plasma membrane fragments containing CD9 were being transferred from OVCAR4 cells they were labelled with PKH67, a green fluorescent lipophilic membrane dye, before coculture with NK-92 cells. After 24h, cells were stained with antibodies against CD45 (for gating NK-92 cells) and CD9 and processed for fluorescence-based flow cytometry (STAR Methods). At the highest ratios of OVCAR4 to NK-92 cells, PKH67 and CD9 were co-detected in ~50% NK-92 cells. As these cell ratios decreased so did the capture of OVCAR4 membrane fragments by NK-92 cells (Figure 5E).

We performed microscopy to visualize CD9 contained within the membrane fragments that were transferred from OVCAR4 to NK-92 cells (Figure 5F). Each cell line was labelled with a lipophilic fluorescent dye; OVCAR4 with PKH67 (green) and NK-92 with PKH26 (red). After coculture for 3h, cells were stained with antibodies against CD9 (blue) and CD45 (white). OVCAR4 cells were only visible in the green (PKH67) and blue (CD9) detection channels. A proportion of NK-92 cells (staining red and positive for CD45) were visible in all four channels confirming their acquisition of OVCAR4 plasma membrane (green) expressing CD9+ (blue). These images, together with the merged channel images demonstrate that NK-92 can trogocytose plasma membrane fragments containing anchored CD9 from OVCAR4 cells (Figure 5F).

### Evaluation of NK-92 trogocytosis from non-HGSC cell lines

In order to determine whether NK-92 trogocytosis of plasma membrane containing CD9 was also a feature with non-HGSC tumor cells, we investigated the frequency of CD9-expressing cells in 15 non-HGSC tumor cell lines (as well as the 11 HGSC tumor cell lines discussed above) (Figure 6A). Since Daubeuf et al. showed that the expression level of a plasma membrane protein was not a determinant of how well it was transferred to the recipient cell we chose three non-HGSC tumor cell lines with high CD9 expression (HCT116, A431 and MCF7) and three with lower CD9 expression (HeLa, CaCo-2 and HepG2) for trogocytosis experiments (Figure 6B) (Daubeuf et al., 2010a). NK-92-mediated trogocytosis from these cell lines was highly varied and confirmed Daubeuf’s observation that it was not correlated to levels of CD9 expression in the effector cells. Apart from the colorectal cancer cell line HCT116, the process was less pronounced than that seen with HGSC cell lines (Figure 5C and 6C). Cytochalasin D inhibition was most marked for the HCT116, MCF7 and CaCo2 cell lines (Figure 6D).

### CD9+ NK-92 cells have a more immune-suppressive intracellular cytokine profile than their CD9-counterpart

It is recognized that d-NK cells exert immune tolerance through their poor cytotoxic responses and a cytokine secretion profile that promotes tolerance and placental development (Crespo et al., 2017; Hanna et al., 2006; Jabrane-Ferrat, 2019). Thus, we hypothesized that on acquiring CD9, NK-92 cells would exhibit similar functions of immune tolerance. To this end we assembled a CyTOF antibody panel designed to measure degranulation, production of cytolytic proteins and intracellular cytokine production (ICP) (Bryceson et al., 2010; Siebert et al., 2008) (Table S6). We analyzed NK-92 cells in monoculture, after coculture with HGSC cell lines and in the presence and absence of phorbol-12-myristate-13-acetate (PMA). Exposing cells to PMA is a convenient method for measuring NK cell function since it bypasses upstream signaling through NK cell receptors (Shabrish et al., 2016) (STAR Methods). The K562 erythroleukemic cell line which only expresses ligands for activating NK receptors was used as a sensitive control for NK cell functional activation (Tremblay-McLean et al., 2019). Inclusion of a CD9 antibody in the CyTOF panel enabled us to gate out CD9+ from CD9-NK-92 cells. For intracellular proteins two metrics were measured: i) changes in frequency of NK-92 cells expressing a specific cytokine (Figure 7A) and ii) median expression of intracellular cytokine production Figure 7B).

**Figure 7:**
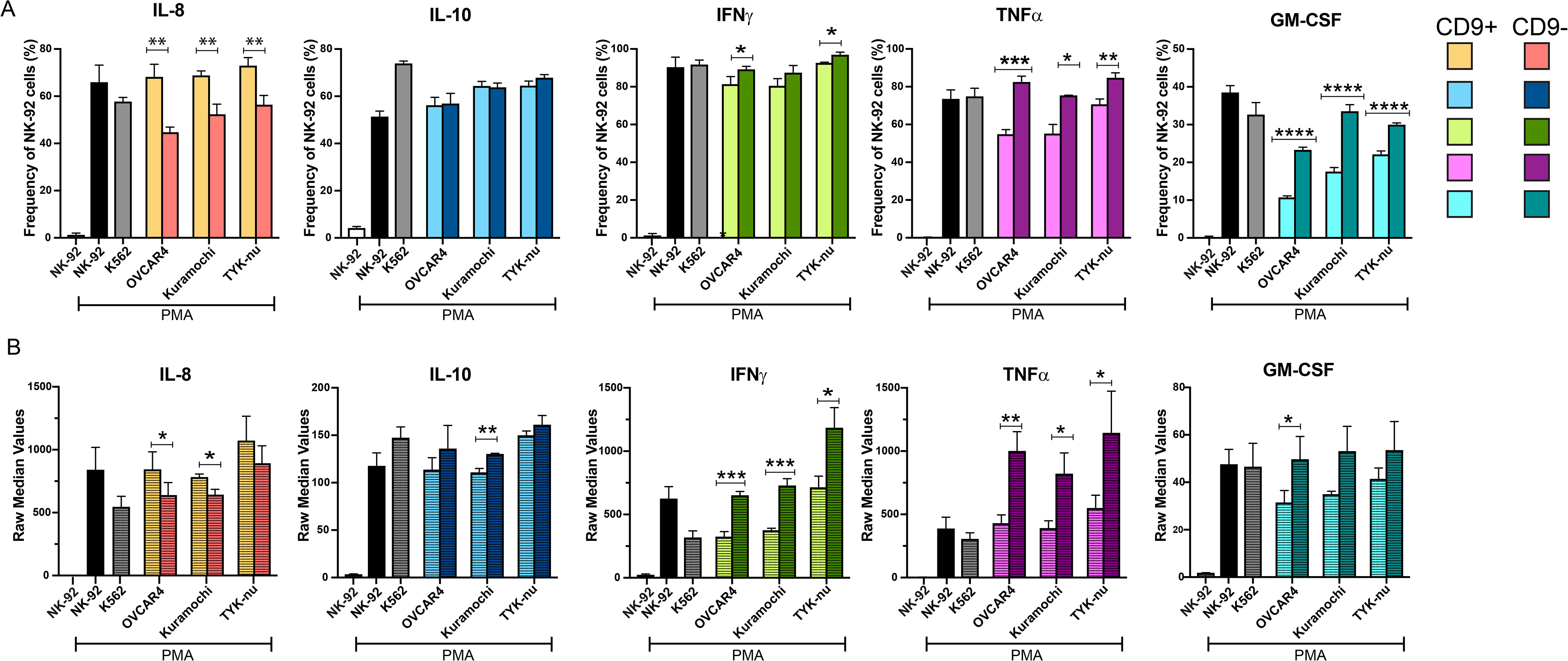
Intracellular cytokine production by CD9+ and CD9-NK-92 cells. HGSC and NK-92 cells (1:1) were cocultured for 6h, treated with PMA/ionomycin or vehicle control and brefeldin A/monensin and processed for CyTOF with the NK cell antibody panel (Table S5 and STAR Methods). CD9+ and CD9-cells were manually gated from the CD45+ cell population. **(A)** Frequency of CD9+ and CD9-cells producing each cytokine as indicated. **(B)** Levels (raw median counts) of cytokine produced by CD9+ and CD9-NK-92 cells. Plots show means of triplicates with standard deviations. Student’s two-tailed t-tests determined statistically significant functional differences between CD9+ and CD9-NK-92 cells. p-values: * ≤ 0.01, ** ≤ 0.001, *** ≤ 0.0001, **** ≤ 0.00001. See also Figure S7 and Table S6.

For a subset of proteins, no differences were observed between NK-92 cells grown in mono-or coculture with or without PMA. High levels of granzyme B, perforin and MIP1β were produced by >85% of NK-92 cells under all conditions with no differences in CD9+ or CD9-NK-92 cells (Bryceson et al., 2010; Fauriat et al., 2010; Siebert et al., 2008). However, cell frequencies and median expression levels of CD107a, a marker for degranulation, and production of MIP1α, were both increased in response to PMA but no differences were seen between CD9+ and CD9-NK-92 cells (Alter et al., 2004). VEGF levels were constitutively high with > 90% of NK-92 cells producing this angiogenic factor in monoculture and after coculture (Figure S7).

After coculture with all three HGSC cell lines and with PMA stimulation, both the frequency and amounts of IL-8 were statistically greater in CD9+ NK-92 cells compared to CD9-NK-92 cells (Figure 7A, B). IL-8 is a proangiogenic factor produced by d-NK cells with a role in vascularizing the placenta (Jabrane-Ferrat, 2019) and in the context of malignancy has a role in promoting the tumor angiogenic system (Yoneda et al., 1998). By contrast, after coculture and PMA stimulation, both cell frequencies and amounts of the anti-tumor cytokines, TNFα, GM-CSF and IFNγ, were statistically lower in CD9+ NK-92 compared to CD9-cells (Bryceson et al., 2010; Siebert et al., 2008) (Figure 7A, B). Similar frequencies (~60%) of CD9+ and CD9-NK-92 cells produced the immunosuppressive cytokine IL-10, but expression levels were reduced in CD9+NK-92 cells after coculture with the Kuramochi cell line (Fauriat et al., 2010) (Figure 7A). These ICP data indicate that NK-92 cells that acquire CD9 are functionally more immune-tolerant.

### HGSC cell lines are poor targets for NK-92 mediated cytotoxicity

NK-92 cells were cocultured with OVCAR4, Kuramochi and TYK-nu cell lines and cytotoxicity was determined by the calcein release assay (Neri et al., 2001) (Figure 8A, STAR Methods). Compared with the K562 cell line, NK-92 cytotoxicity was significantly reduced toward all three HGSC cell lines. Furthermore, the magnitude of attenuation trended with stage of tumor progression such that OVCAR4 cells were more, and TYK-nu cells less susceptible to NK-92 cell-mediated killing consistent with the more immunosuppressive NK receptor ligand profile for V cells described above (Figures 2, 3, 8A).

**Figure 8.**
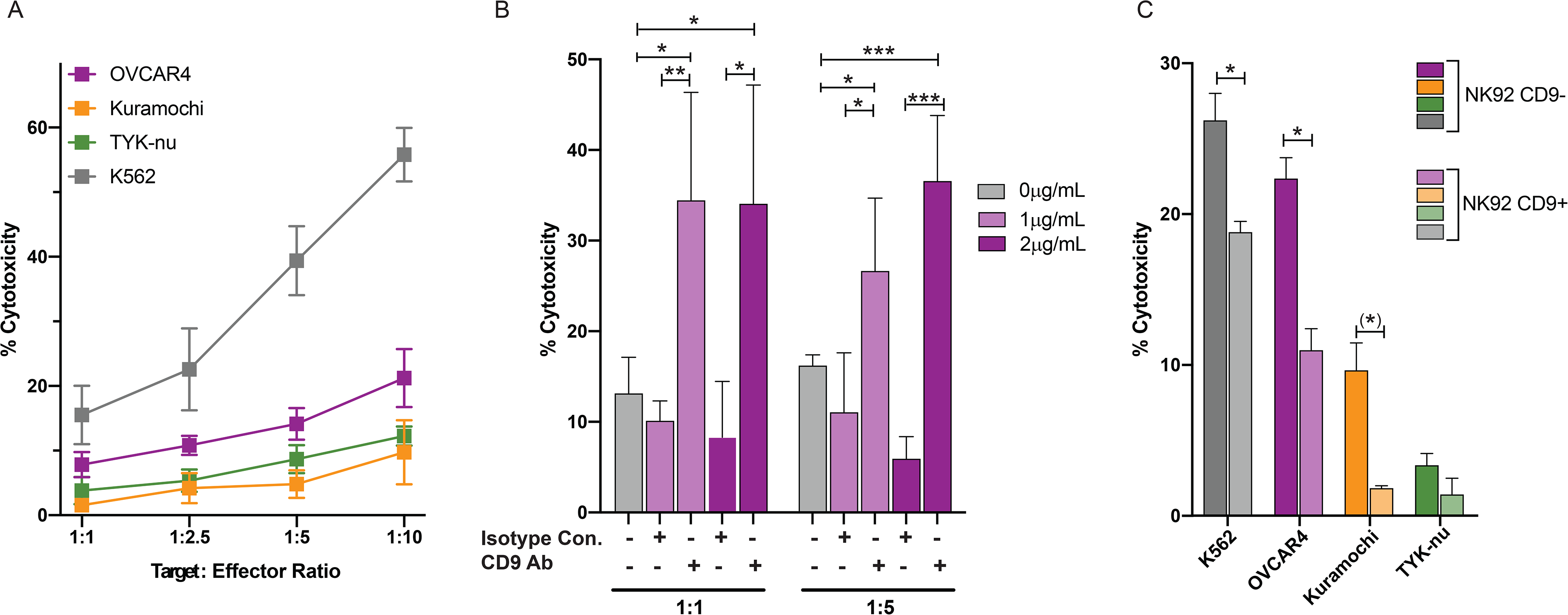
NK-92-mediated cell cytotoxicity toward HGSC cell lines. Coculture HGSC cell lines (as indicated) were assayed for their susceptibility to NK-92 mediated cytotoxicity by the calcein release assay. **(A)** NK-92 cells have reduced cytotoxicity toward HGSC cell lines compared to the control K562 cell line, at the target: effector ratios shown. **(B)** Addition of a CD9 blocking antibody significantly increased NK-92 cytotoxicity. Data are shown for quadruplets performed with two antibody concentrations and different target : effector cell ratios. Statistical significance determined with two tailed t test: * p ≤ 0.05, ** p ≤ 0.01, p ≤ 0.001. **(C)** FACS-sorted CD9+ NK-92 after coculture have a reduced cytotoxicity function compared to CD9-NK-92 cells grown in monoculture. Statistical significance determined with two tailed t test: * p ≤ 0.03, (*) p = 0.06.

### CD9 antibody blockade increases NK-92 cell-mediated cytotoxicity

To determine the contribution of CD9 to the attenuated cytotoxicity of HGSC cells, we performed the calcein release assay in the presence of a CD9 blocking antibody (Figure 8B). The data showed that the CD9 blocking antibody significantly increased NK-92-mediated cytotoxicity for OVCAR4 cells. These data make a strong case for CD9 having a prominent immunosuppressive role.

### Purified CD9+ NK-92 cells exhibit reduced cytotoxicity

To directly compare cytotoxicity of CD9+ with CD9-NK-92 cells, we FACS-sorted CD9+ NK-92 cells after coculture with OVCAR4. The calcein release cytotoxicity assay revealed significant attenuation of killing activity by CD9+ after coculture with OVCAR4 and Kuramochi cells compared to CD9-NK-92. The TYK-nu cell line was very resilient to both CD9+ and CD9-NK-92 cell-mediated killing consistent with its mesenchymal phenotype (Figure 8C) (Gonzalez et al., 2018). These data demonstrate that the presence of CD9 on NK-92 cells endows them with a more immune-tolerant phenotype.

## Discussion

This study was designed to identify T and NK cell types within the tumor infiltrate of newly diagnosed HGSC that could provide insight into the disappointing results from clinical trials with immune checkpoint blockade treatments.(Kandalaft et al., 2019). In line with this, our CyTOF data analysis of newly diagnosed, chemo-naïve HGSC tumors revealed previously unidentified dl-NK cell subpopulations that were positively correlated with the overall abundance of tumor and EV cells (Figure 1A - C) (Koopman et al., 2003; Kopcow et al., 2005). The presence of dl-NK cells has been reported in colorectal and lung tumors, but to our knowledge this is the first report of this immune cell type in HGSC (Bruno et al., 2018).

Decidual NK cells express the tetraspanin CD9 and are functionally distinct from peripheral NK cells (Jabrane-Ferrat, 2019; Koopman et al., 2003). They comprise 70% of the total lymphocyte population during the first trimester of pregnancy and produce a wide range of secretory proteins that are critical for decidualization, formation and vascularization of the placenta and crucially, creation of a privileged immune tolerant maternal-fetal compartment (Hanna et al., 2006; Jabrane-Ferrat, 2019; Venkitaraman, 2009). Furthermore, although d-NK cells are poorly cytotoxic they contain cytotoxic granules which can be transiently activated to provide immunity to infection during pregnancy (Jabrane-Ferrat, 2019; Tilburgs et al., 2015; Vento-Tormo et al., 2018).

This study was based on the recognition that HGSC tumor cells promote immune tolerance to support tumor expansion. The identification and characterization of dl-NK cells described here supports the notion that HGSC tumor cells have a role in modulating NK cells toward a dl-NK phenotype. This included transfer of the tetraspanin CD9, a marker that classifies d-NK cells, by trogocytosis as well as co-expression of NK receptor ligands that increase immunosuppression of NK cells.

We previously reported on CyTOF analysis of the tumor compartment in newly diagnosed HGSC and have now extended our analysis to include the immune infiltrates of these tumors and to investigate how the tumor cells interact with the immune cells therein. We simultaneously measured the expression of twelve NK receptor ligands and two ADAMs proteases by the tumor cells. This contrasts with other correlative studies that measured only one or two of these ligands (Lopez-Soto et al., 2017; Lorenzo-Herrero et al., 2018). CyTOF data revealed that E and EV tumor cells expressed more NK activating receptor ligands than V cells consistent with previous reports that described loss of NKG2D ligands in tumor cells undergoing epithelial to mesenchymal transition (Huergo-Zapico et al., 2018; Lopez-Soto et al., 2017) (Figures 2 and 3). The greater number of ligand combinations in earlier stages of tumor development could increase the likelihood of tumor cell escape from immune surveillance with eventual transition into poor-prognosis V cells. Having attained a V phenotype, these tumor cells may then switch to alternative more stringent mechanisms of immune escape to promote their metastasis (Figures 2D and 3). Additionally, high levels of ULBP4, a functional outlier within the ULBP family, were observed in V cells and has been shown to promote immune escape by a variety of mechanisms (Dhar and Wu, 2018; Labani-Motlagh et al., 2016; Zoller et al., 2018) (Figure 2B).

Numerous studies documented upregulation of NK receptor activating ligands in response to exposure to genotoxic agents (Dhar and Wu, 2018; Gasser and Raulet, 2006), epigenetic modifiers (Bugide et al., 2018) and radiation (Gasser et al., 2005). Thus, carboplatin used to treat patients with HGSC, could also upregulate these ligands and potentially make HGSC more amenable to NK immunotherapy (Bowtell et al., 2015; Matulonis, 2018). However, our results showed that carboplatin induced a more inhibitory NK receptor ligand phenotype (increased HLA-E, nectin 4 and HLA-ABC) in E, EV and V HGSC cell lines, revealing an unrecognized mechanism of carboplatin resistance (Figure 4). Nectin 4 was recently shown to be a ligand for the inhibitory receptor TIGIT and, in a separate study nectin 4 expression was proposed to have a role in HGSC metastasis and chemotherapeutic resistance (Derycke et al., 2010; Ito et al., 2018; Reches et al., 2020). Furthermore, both nectin 4 and CD111 have additional roles in adhesion, cell movement and stem cell biology (Belaaloui et al., 2003; Martinet and Smyth, 2015; Yurtsever et al., 2013). Characterizing NK receptor ligand expression, especially after carboplatin-based chemotherapy may be valuable for determining whether a patient is a candidate for NK immunotherapy.

To gain functional insight into how HGSC cells might modulate NK cells towards an immunosuppressive role we selected E, EV and V HGSC cell lines for coculture with NK-92 cells (Domcke et al., 2013). The NK-92 cell line expresses CD56, lacks expression of inhibitory KIR receptors and is molecularly well-characterized. NK-92 cells that are unmodified or genetically engineered to express chimeric antigen receptors or activating NK receptors such as NKG2D, have satisfied safety criteria in several early phase clinical trials (Daher and Rezvani, 2018; Rezvani, 2019; Rezvani et al., 2017; Suck et al., 2016).

Our data confirmed that CD9 protein was transferred from HGSC cells to NK-92 cells by NK-92-mediated trogocytosis. (Figure 5). This indicates the potential ease with which NK cells could be reprogrammed toward a dl-NK cell phenotype. (Figure 5). Trogocytosis has been observed for T cells, B cells, basophils and NK cells (Domaica et al., 2009; Gary et al., 2012; Joly and Hudrisier, 2003; McCann et al., 2007; Miyake et al., 2017; Quah et al., 2008; Tabiasco et al., 2002). In vitro coculture experiments between d-NK cells and extravillous trophoblasts, and in a separate study, between peripheral NK and melanoma cells resulted in the transfer of HLA-G onto the NK cells and enhanced their immune tolerance.(Caumartin et al., 2007; Tilburgs et al., 2015). Additionally, a recent study using a mouse model of B-cell acute lymphoblastic leukemia described trogocytosis as the mechanism by which chimeric antigen receptor T (CAR-T) cells acquired the CD19 target they were engineered to seek out from the tumor cells they were programmed to kill (Hamieh et al., 2019). This mechanism of therapeutic resistance through trogocytosis could account for the observed tumor cell antigen loss in patients undergoing CAR-T cell therapy (Hamieh et al., 2019). Furthermore, the tetraspanin CD81, which is complexed with CD19, was jointly acquired and raises the question as to whether tetraspanins have a central role in trogocytosis (Daubeuf et al., 2010a; Daubeuf et al., 2010b). These studies provide ever-increasing evidence for trogocytosis playing a key role in immune tolerance and therapeutic resistance.

While the exclusivity of CD9 expression on the decidual NK cell subpopulation is well established, its functional role in these cells is unclear (Koopman et al., 2003; Kopcow et al., 2005). In this study, our coculture data analysis demonstrated that gain of CD9 expression by NK-92 cells coincided with decreased production of IFNγ, TNFα and GM-CSF, increased production of IL-8 and suppressed cytotoxicity (Figure 7 and 8). Critically, a CD9 blocking antibody significantly increased NK-92 cytotoxicity providing strong evidence that CD9 confers NK-92 cells with immunosuppressive properties (Figure 8B).

In additional coculture experiments NK-92 cells also acquired CD9 from non-HGSC cell lines but often to greatly diminished extents when compared with OVCAR4 cells (Figure 6C, D). Based on these data, trogocytosis of CD9 is likely to occur in non-HGSC tumors, but at very different frequencies. It is conceivable that dl-NK cells reported in colorectal and lung cancer acquire CD9 by trogocytosis (Bruno et al., 2018).

We observed high levels of CD9 in HGSC tumor cells suggesting that in vivo these tumor cells could be a likely source of CD9, not only for intra-tumoral NK cell-mediated trogocytosis, but potentially for other tumor-infiltrating immune cell types (Figure S5). Thus, we observed two T cell clusters expressing high levels of CD9 that correlated with total tumor and EV cell abundance (Figure S5, Figure S2A). Additionally, two other CD9+ immune cell clusters (CD3-CD56^Lo^ CD9+ CXCR3+ CD4+) showed no correlation with any tumor features but were phenotypically related to both dl-NK and T cell clusters (Figures S2B and S3). Their presence at high frequency in all tumors, suggests they could act as precursors for the correlating dl-NK and T cells and signify a previously unappreciated plasticity between these NK and T-cell phenotypes.

CD9 shows ubiquitous distribution and is involved in multiple cellular functions such as proliferation, motility and adhesion with major roles in formation of the immune cell synapse (Reyes et al., 2018). It is thus likely that CD9 may have multiple roles in regulating the HGSC tumor immune microenvironment. It has been shown to directly associate with ADAM17 protease and inhibit its cleavage activity toward surface protein ectodomains (Gutierrez-Lopez et al., 2011). Thus, transfer of CD9 from HGSC tumor cells onto NK-92 cells could reactivate ADAM17 to cleave NK activating receptor ligand substrates thereby indirectly facilitating immune escape by ligand and/or receptor shedding (Boutet et al., 2009; Ferrari de Andrade et al., 2018; Lanier, 2015; Okumura et al., 2020; Raulet et al., 2013). The precise mechanism by which CD9, or other co-transferred proteins, suppressed NK-92 immune cell function remains to be established.

The ever-increasing number and complexity of tumor-immune escape mechanisms for HGSC are particularly evident in a recent study showing the presence of distinct inter-regional tumor-immune subtypes within and between HGSC patients (Zhang et al., 2018). This further highlights the challenges associated with choosing predictive biomarkers to guide the choice of the most beneficial immunotherapy. One invaluable source of information in the foreseeable future would be from the analysis of tumor samples from HGSC patients who are long-term survivors (Garsed et al., 2018).

The significance of our results underscores the critical need to evaluate both CD9 and NK receptor ligand expression within HGSC tumors to stratify those patients most likely to benefit from NK immunotherapy. Furthermore, the abundant expression of CD9 on HGSC tumor cells presents the distinct possibility that NK-92 or other adoptively transferred NK cells could readily acquire CD9 by trogocytosis and transition to a more immunosuppressive phenotype which could negatively affect administered immunotherapy (Hermanson et al., 2016; Rezvani, 2019; Suck et al., 2016). Thus, one logical next step would be to determine the feasibility of a peripheral blood test to monitor gain of CD9 by adoptively transferred NK cells.

The data from this study identify previously unrecognized mechanisms of immune suppression in HGSC and support the development of CD9 as a potential drug target with immediate relevance for NK immunotherapy.

## Acknowledgements

The authors wish to thank Drs. Catherine Blish and Matija Peterlin for critical reading of the manuscript. We thank Drs. Catherine Blish and Nancy Zhao for help with the calcein release assay.

## Author contributions

V.D.G designed and performed experiments, analyzed, and interpreted data and wrote manuscript. Y-W.H designed and performed experiments, analyzed, and interpreted data. S-Y.C designed and performed experiments, analyzed, and interpreted data. A. D-G performed coculture experiments and analyzed data. K.D performed microscopy. K.S and A. G analyzed and interpreted data. E.P designed and performed experiments, analyzed data, and wrote manuscript. W.J.F designed experiments, analyzed, and interpreted data, wrote manuscript, and supervised the work.

## STAR Methods

### Patient Samples

Deidentified newly diagnosed chemo naïve HGSC tumors prepared as single cell suspensions for CyTOF analysis collected over a two-year period were purchased from Indivumed (Hamburg, Germany) (Table S2). Tumor samples were collected in compliance with the Helsinki declaration and all patients provided written informed consent. The use of human tissue was approved and complied with data protection for patient confidentiality. Institutional review board approval was obtained at the Physicians Association in Hamburg, Germany.

### Genomic sequencing and analysis for TP53 and BRCA1/2

DNA was extracted and enriched through multiplex PCR (QIAGEN QIAmp DNA Mini-Kit and QIAGEN GeneRead DNaseq Targeted Ovarian V2 Panel, respectively). The TrueSeq protocol was used to make an indexed Illumina sequencing library from the pooled sample amplicons. The subsequent protocols for sequencing were described previously(Gonzalez et al., 2018). The pathogenic variants were noted (Table S3).

### Cancer cell lines from HGSC and non-HGSC malignancies

Cell lines were authenticated by short tandem repeat (STR) profiling performed by the Stanford Functional Genomics Facility. Ovarian cancer cell lines (OVCAR4, Kuramochi, and TYK-nu), NK-92 and other non-HGSC cell lines used for CD9 screen were grown according to the recommended conditions from their respective vendors (Table S7).

### Antibodies for CyTOF

Antibodies were either purchased pre-conjugated or conjugated in-house as previously reported(Gonzalez et al., 2018) In brief, for in-house conjugations, antibodies in carrier-free PBS were conjugated to metal-chelated polymers (MaxPAR antibody conjugation kit, Fluidigm) according to the manufacturer’s protocol or to bismuth with our protocol(Han et al., 2017). Metal-labeled antibodies were diluted to 0.2–0.4 mg/mL in antibody stabilization solution (CANDOR Biosciences) and stored at 4°C. Each antibody was titrated using cell lines and primary human samples as positive and negative controls. Antibody concentrations used in experiments were based on an optimal signal-to-noise ratio. Three CyTOF antibody panels were used in this study to characterize: i) tumor T and NK cells (Table S3) ii) Tumor NK receptor ligand expression and (Table S4) iii) NK cell receptor and intracellular cytokine expression (Table S6).

### Antibodies for Fluorescence-Based Flow Cytometry

Antibodies were purchased for detection of CD9 and CD45 from Becton Dickinson (CD9 BV421, CD9 PE) and Biolegend (CD45 APC). The same antibody clones were used for CyTOF (Tables S3, S4 and S6). Near-IR fixable LIVE/DEAD stain from Thermo Fisher Scientific was used to distinguish dead cells.

### Sample Processing and Antibody Staining for CyTOF

Frozen, fixed single-cell suspensions of HGSC tumors or cell lines were thawed at room temperature. For each sample, 1 × 10^6^ cells were aliquoted into cluster tubes in 96 well plates and subjected to pre-permeabilization palladium barcoding(Gonzalez et al., 2018). After barcoding, pooled cells were pelleted and incubated with Trustain FcX Fc receptor block (Biolegend) to prevent non-specific antibody binding, for 10min at room temperature. Cells were then incubated with antibodies against surface markers for 45min at room temperature. Cells were permeabilized at 4°C with methanol or 1x Permeabilization Buffer (eBioscience, Thermo Fisher Scientific), (only for NK cell CyTOF antibody panel Table S6) on ice for 10min. Cells were subsequently stained with antibodies against intracellular markers for 1h at room temperature, washed, and incubated with the ^191/193^Ir DNA intercalator (Fluidigm) at 4°C overnight. Cells were washed and resuspended in a solution of normalization beads before introduction into the CyTOF 2 (Bendall et al., 2011; Gonzalez et al., 2018).

### Determination of intracellular pools of CD9 by CyTOF

NK-92 and OVCAR4 cells were stained with cisplatin (Sigma Aldrich), fixed with 1.6% paraformaldehyde (Thermo Fisher Scientific), washed, incubated with Trustain FcX Fc receptor block (Biolegend) (as above) and then incubated with CD9-PE (Becton Dickinson) for 45 min at room temperature. Cells were washed and stained with anti-PE-165Ho (Fluidigm) for 30min at room temperature. Following secondary antibody staining, cells were permeabilized with 1x Permeabilization Buffer ((eBioscience, Thermo Fisher Scientific), on ice for 10min. Cells were subsequently stained with CD9-156Gd (Fluidigm) (to detect intracellular CD9) for 1h at room temperature. Cells were processed and introduced into the CyTOF 2 as described above.

### In vitro cocultures to determine intracellular cytokine production of NK-92 cells

The HGSC cell lines, OVCAR4, Kuramochi and TYK-nu cells, were each cocultured with the NK-92 cell line, at an effector:target ratio of 1:1 for 6h unless otherwise indicated, at 37°C in a humidified cell culture incubator. HGSC cells (100,000 cells / well) were seeded in U-bottom 96-well plates (Corning, Costar) with NK-92 cells (100,000). During the last 4h of coculture, PMA / Ionomycin cell stimulation cocktail (500X) ((eBioscience, Thermo Fisher Scientific), was added to induce intracellular cytokine production. The protein transport inhibitors, Brefeldin A and Monensin (eBioscience, Thermo Fisher Scientific), were used at a final concentration of 3μg/ml and 2μM, respectively. There were two positive controls; i) NK-92 cells grown in monoculture −/+ PMA and ii) coculture of the K562 cell line (HLA-null erythroleukemic) (Tremblay-McLean et al., 2019) with NK-92 cells. CD107a-151Eu antibody (Fluidigm) (1μl) was added to each well as a marker for degranulation. All experiments were performed with biological and technical triplicates with details described for specific assays.

### Transwell assay

OVCAR4, Kuramochi and TYK-nu cells were cocultured with NK-92 in a 96-well dual-chamber transwell plate with 3μm micropores (Corning, Costar). HGSC cells (100,000/well) were placed into the lower chamber, and NK-92 cells (100,000/well) were placed into the upper chamber. The cells were cultured at 37°C for 6h in a humidified cell culture incubator. The assay was performed with biological and technical triplicates.

### Trogocytosis

OVCAR4 were labeled with PKH67 (Sigma Aldrich) prior to coculture with NK-92. In brief, OVCAR4 were washed with serum free media and resuspended in diluent C. A 2X working solution of PKH67 was prepared immediately prior to use. Cells were mixed with PKH67 working solution for a final concentration of 5×10^6^ cells/mL in 20μM PKH67 and incubated for 5min at room temperature. The labeling was quenched with an equal volume of fetal bovine serum, incubated for 1min, and washed three times with 10mL of complete media. Cells were seeded in U-bottom 96-well plates (Corning, Costar), and cocultured with NK-92 cells at target : effector ratios 5:1, 2.5:1, 1:1, 2.5:1 and 5:1 for 24h at 37°C. Cells were then stained with CD45 and CD9 antibodies and processed for flow cytometry.

### Reverse transcriptase quantitative PCR to measure CD9 transcript levels

Total mRNA was isolated with a Qiagen MicroRNA isolation kit from OVCAR4 and NK-92 cells grown in monoculture and FACS-sorted CD9+ and CD9-NK-92 cells after coculture with OVCAR4 cells. cDNA was generated using the High Capacity cDNA Reverse Transcription Kit from Applied Biosystems according to manufacturer’s protocol. Real-time PCR for E-cadherin (CDH1) Hs00170423_m1, CD45 (PTPRC) Hs00894716_m1, and CD9 Hs01124022 was performed with a Taqman gene expression kit and run on the ABI 7900HT instrument.

### Microscopy to image trogocytosis

OVCAR4 and NK-92 cells were labeled with the membrane dyes PKH67 and PKH26 (Sigma Aldrich), respectively, prior to coculture. Cells were washed with serum free media and resuspended in Diluent C. A 2X working solution of each membrane dyes was prepared immediately prior to use. Cells (5×10^6^cells/mL) were mixed with their respective working solution of dye for a final concentration of 20μM. After a 5min incubation at room temperature the labeling was quenched with an equal volume of fetal bovine serum, incubated for 1min, and washed three times with 10mL of complete media. Cells were seeded in U-bottom 96-well plates (Corning, Costar), and cocultured with NK-92 cells at an effector:target ratio 1:1 for 3h at 37°C. Cells were then fixed with a final concentration of 1.6% paraformaldehyde, stained with CD45 and CD9 antibodies and seeded on microscope slides for imaging on a Keyence BZ-X800 microscope.

### Trogocytosis inhibition

NK-92 cells were pre-treated with cytochalasin D (10μM in complete media) for 2h (Aucher et al., 2008). They were then co-cultured with cancer cell lines as indicated in a ratio of 1:1 in the continuous presence of cytochalasin D for a further 2h after which cells were stained with antibodies against CD9, CD45 and processed prior to CyTOF analysis as described above.

### Calcein-AM release cytotoxicity assay

OVCAR4, Kuramochi, TYK-nu and K562 (control) target cells were washed in PBS and resuspended in calcein-acetoxymethyl (calcein-AM; Thermo Fisher Scientific) staining solution (2.5μM in PBS) at a cell density of 1 × 10^6^/mL and incubated for 30min at 37°C (Lorenzo-Herrero et al., 2020; Ramadoss et al., 2019). Target cells were seeded in U-bottom 96-well plates (Corning, Costar), and cocultured with NK-92 cells at increasing effector : target ratios 1:1, 2.5:1, 5:1 and 10:1 for 4 h at 37°C. For the cytotoxicity assay with CD9 blocking antibody, after both cells lines were plated either control mouse IgG1 kappa (Abcam, clone: 15-6E10A7) or purified mouse monoclonal CD9 antibody (Abcam, clone: MEM-61) were added to the coculture at the concentrations indicated for the duration of the incubation. Cells were then spun down and 100μl of supernatant were transferred to a black-walled 96 well plate (Corning, Costar). Calcein release was measured from the fluorescent signal using 485 nm excitation wavelength and 530 nm emission wavelength (Ex/Em Calcein: 494/517 with a Tecan Infinite M1000 fluorescent plate reader. Control wells contained HGSC target cells alone (spontaneous lysis) or with 2% Tween-20 (maximum lysis). Specific killing was calculated using the equation: specific killing = (lysis of coculture - spontaneous lysis) / (maximum lysis - spontaneous lysis) x 100%. The assay was performed with biological and technical quadruplets.

### Data analysis tools

All data and statistical analysis were implemented with Microsoft Excel, Matlab, R and GraphPad Prism 8. CyTOF datasets were evaluated with software available from Cytobank(Kotecha et al., 2010) and CellEngine (cellengine.com, Primity Bio).

### Initial Assessment of data quality

Initial data quality was assessed by determining dead and apoptotic cells which were excluded from further analysis. Viable cells, defined as cisplatin negative and cleaved PARP negative were used for experiments (Gonzalez et al., 2018). For experiments with newly diagnosed HGSC tumors, tumor cells were gated as CD45-/CD31-/FAP- and immune cells were gated as CD45+ CD66-as described previously (Gonzalez et al., 2018).

### Clustering of tumor and T and NK cell immune infiltrate

Manually gated CD45+CD66-(immune) and CD45-/CD31-/FAP-(tumor) cells from each HGSC tumor were pooled for clustering with X-shift (Samusik et al., 2016), a density-based clustering algorithm, using the Vortex clustering environment (http://web.stanford.edu/_samusik/vortex). Markers for clustering the immune and tumor cells are shown in Tables S3 and S4, respectively.

### Correlation network analysis

Spearman pairwise correlation coefficients (r_s_) were computed between: i) cell frequencies for 56 tumor cell clusters, ii) frequency of 52 T and NK cell clusters, iii) total tumor cell abundance, iv) total EV cell abundance, v) total E-cadherin cell abundance and vi) other features previously described (Gonzalez et al., 2018). A hierarchically ordered heat-map was generated in R.

### Force directed layout visualization

Force directed layouts were generated from a composite of all 12 HGSC tumors. After merging all single cell data files, cells were clustered. 10,000 single cells were computationally sampled from each of the 56 tumor cell clusters. Each cell was connected on a 10-nearest-neighbor graph. This graph was subjected to a force-directed layout (FDL) that placed groups of phenotypically related cells adjacent to one another (Samusik et al., 2016). Repeat samplings generated comparable results. Layouts were colored for expression of E-cadherin, vimentin, NK receptor ligands or ADAM10 and ADAM17.

### Combinatorial expression for NK receptor ligands in E, EV and V tumor compartments

The tumor population CD45-/FAP-/CD31-was the parent population for gating the E, EV and V tumor compartments (Gonzalez et al., 2018). Frequencies of tumor cell subpopulations defined by their combinatorial expression patterns of the twelve NK receptor ligands and two ADAM proteases were determined with MATLAB. For this analysis, the frequency of tumor cells expressing each of these proteins was determined in each compartment on a per sample basis. Combinations used in the analysis were based on a threshold frequency of >1% for cells in any compartment in any sample.

### Simpson’s index of diversity

The Simpson’s index of diversity, D was calculated in Excel with the formula 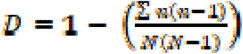 where N is the total number of tumor cell subpopulations with a specific NK receptor ligand combination (163) and n is the number of times a subpopulation is present in the E, EV and V compartments of each of the 12 tumors.

## References

Alter, G., Malenfant, J.M., and Altfeld, M. (2004). CD107a as a functional marker for the identification of natural killer cell activity. Journal of immunological methods 294, 15–22.

Ashworth, A., and Lord, C.J. (2018). Synthetic lethal therapies for cancer: what’s next after PARP inhibitors? Nat Rev Clin Oncol 15, 564–576.

Aucher, A., Magdeleine, E., Joly, E., and Hudrisier, D. (2008). Capture of plasma membrane fragments from target cells by trogocytosis requires signaling in T cells but not in B cells. Blood 111, 5621–5628.

Bast, R.C., Jr., Matulonis, U.A., Sood, A.K., Ahmed, A.A., Amobi, A.E., Balkwill, F.R., Wielgos-Bonvallet, M., Bowtell, D.D.L., Brenton, J.D., Brugge, J.S., and Coleman, R.L. (2019). Critical Questions in Ovarian Cancer Research and Treatment: Report of an American Association for Cancer Research Special Conference.

Belaaloui, G., Imbert, A.M., Bardin, F., Tonnelle, C., Dubreuil, P., Lopez, M., and Chabannon, C. (2003). Functional characterization of human CD34+ cells that express low or high levels of the membrane antigen CD111 (nectin 1). Leukemia 17, 1137–1145.

Bendall, S.C., Simonds, E.F., Qiu, P., Amir el, A.D., Krutzik, P.O., Finck, R., Bruggner, R.V., Melamed, R., Trejo, A., Ornatsky, O.I., et al. (2011). Single-cell mass cytometry of differential immune and drug responses across a human hematopoietic continuum. Science 332, 687–696.

Bernstein, H.B., Plasterer, M.C., Schiff, S.E., Kitchen, C.M., Kitchen, S., and Zack, J.A. (2006). CD4 expression on activated NK cells: ligation of CD4 induces cytokine expression and cell migration. J Immunol 177, 3669–3676.

Boutet, P., Aguera-Gonzalez, S., Atkinson, S., Pennington, C.J., Edwards, D.R., Murphy, G., Reyburn, H.T., and Vales-Gomez, M. (2009). Cutting edge: the metalloproteinase ADAM17/TNF-alpha-converting enzyme regulates proteolytic shedding of the MHC class I-related chain B protein. J Immunol 182, 49–53.

Bowtell, D.D., Bohm, S., Ahmed, A.A., Aspuria, P.J., Bast, R.C., Jr., Beral, V., Berek, J.S., Birrer, M.J., Blagden, S., Bookman, M.A., et al. (2015). Rethinking ovarian cancer II: reducing mortality from high-grade serous ovarian cancer. Nat Rev Cancer 15, 668–679.

Brennick, C.A., George, M.M., Corwin, W.L., Srivastava, P.K., and Ebrahimi-Nik, H. (2017). Neoepitopes as cancer immunotherapy targets: key challenges and opportunities. Immunotherapy 9, 361–371.

Bruno, A., Bassani, B., D’Urso, D.G., Pitaku, I., Cassinotti, E., Pelosi, G., Boni, L., Dominioni, L., Noonan, D.M., Mortara, L., and Albini, A. (2018). Angiogenin and the MMP9-TIMP2 axis are up-regulated in proangiogenic, decidual NK-like cells from patients with colorectal cancer. Faseb j 32, 5365–5377.

Bryceson, Y.T., Fauriat, C., Nunes, J.M., Wood, S.M., Bjorkstrom, N.K., Long, E.O., and Ljunggren, H.G. (2010). Functional analysis of human NK cells by flow cytometry. Methods Mol Biol 612, 335–352.

Bugide, S., Green, M.R., and Wajapeyee, N. (2018). Inhibition of Enhancer of zeste homolog 2 (EZH2) induces natural killer cell-mediated eradication of hepatocellular carcinoma cells. Proc Natl Acad Sci U S A 115, E3509–e3518.

Caumartin, J., Favier, B., Daouya, M., Guillard, C., Moreau, P., Carosella, E.D., and LeMaoult, J. (2007). Trogocytosis-based generation of suppressive NK cells. Embo j 26, 1423–1433.

Cerboni, C., Fionda, C., Soriani, A., Zingoni, A., Doria, M., Cippitelli, M., and Santoni, A. (2014). The DNA Damage Response: A Common Pathway in the Regulation of NKG2D and DNAM-1 Ligand Expression in Normal, Infected, and Cancer Cells. Frontiers in immunology 4, 508.

Chiossone, L., Dumas, P.Y., Vienne, M., and Vivier, E. (2018). Natural killer cells and other innate lymphoid cells in cancer. Nat Rev Immunol 18, 671–688.

Cooper, M.A., Fehniger, T.A., and Caligiuri, M.A. (2001). The biology of human natural killer-cell subsets. Trends in immunology 22, 633–640.

Crespo, A.C., van der Zwan, A., Ramalho-Santos, J., Strominger, J.L., and Tilburgs, T. (2017). Cytotoxic potential of decidual NK cells and CD8+ T cells awakened by infections. J Reprod Immunol 119, 85–90.

Daher, M., and Rezvani, K. (2018). Next generation natural killer cells for cancer immunotherapy: the promise of genetic engineering. Curr Opin Immunol 51, 146–153.

Dance, A. (2019). Core Concept: Cells nibble one another via the under-appreciated process of trogocytosis. Proc Natl Acad Sci U S A 116, 17608–17610.

Daubeuf, S., Aucher, A., Bordier, C., Salles, A., Serre, L., Gaibelet, G., Faye, J.C., Favre, G., Joly, E., and Hudrisier, D. (2010a). Preferential transfer of certain plasma membrane proteins onto T and B cells by trogocytosis. PLoS One 5, e8716.

Daubeuf, S., Lindorfer, M.A., Taylor, R.P., Joly, E., and Hudrisier, D. (2010b). The direction of plasma membrane exchange between lymphocytes and accessory cells by trogocytosis is influenced by the nature of the accessory cell. J Immunol 184, 1897–1908.

Derycke, M.S., Pambuccian, S.E., Gilks, C.B., Kalloger, S.E., Ghidouche, A., Lopez, M., Bliss, R.L., Geller, M.A., Argenta, P.A., Harrington, K.M., and Skubitz, A.P. (2010). Nectin 4 overexpression in ovarian cancer tissues and serum: potential role as a serum biomarker. Am J Clin Pathol 134, 835–845.

Dhar, P., and Wu, J.D. (2018). NKG2D and its ligands in cancer. Curr Opin Immunol 51, 55–61.

Domaica, C.I., Fuertes, M.B., Rossi, L.E., Girart, M.V., Avila, D.E., Rabinovich, G.A., and Zwirner, N.W. (2009). Tumour-experienced T cells promote NK cell activity through trogocytosis of NKG2D and NKp46 ligands. EMBO Rep 10, 908–915.

Domcke, S., Sinha, R., Levine, D.A., Sander, C., and Schultz, N. (2013). Evaluating cell lines as tumour models by comparison of genomic profiles. Nature communications 4, 2126.

Fabre-Lafay, S., Monville, F., Garrido-Urbani, S., Berruyer-Pouyet, C., Ginestier, C., Reymond, N., Finetti, P., Sauvan, R., Adelaide, J., Geneix, J., et al. (2007). Nectin-4 is a new histological and serological tumor associated marker for breast cancer. BMC Cancer 7, 73.

Fauriat, C., Long, E.O., Ljunggren, H.G., and Bryceson, Y.T. (2010). Regulation of human NK-cell cytokine and chemokine production by target cell recognition. Blood 115, 2167–2176.

Ferrari de Andrade, L., Tay, R.E., Pan, D., Luoma, A.M., Ito, Y., Badrinath, S., Tsoucas, D., Franz, B., May, K.F., Jr., Harvey, C.J., et al. (2018). Antibody-mediated inhibition of MICA and MICB shedding promotes NK cell-driven tumor immunity. Science 359, 1537–1542.

Garsed, D.W., Alsop, K., Fereday, S., Emmanuel, C., Kennedy, C.J., Etemadmoghadam, D., Gao, B., Gebski, V., Gares, V., Christie, E.L., et al. (2018). Homologous Recombination DNA Repair Pathway Disruption and Retinoblastoma Protein Loss Are Associated with Exceptional Survival in High-Grade Serous Ovarian Cancer. Clin Cancer Res 24, 569–580.

Gary, R., Voelkl, S., Palmisano, R., Ullrich, E., Bosch, J.J., and Mackensen, A. (2012). Antigen-specific transfer of functional programmed death ligand 1 from human APCs onto CD8+ T cells via trogocytosis. J Immunol 188, 744–752.

Gasser, S., Orsulic, S., Brown, E.J., and Raulet, D.H. (2005). The DNA damage pathway regulates innate immune system ligands of the NKG2D receptor. Nature 436, 1186–1190.

Gasser, S., and Raulet, D.H. (2006). The DNA damage response arouses the immune system. Cancer Res 66, 3959–3962.

Gonzalez, V.D., Samusik, N., Chen, T.J., Savig, E.S., Aghaeepour, N., Quigley, D.A., Huang, Y.W., Giangarra, V., Borowsky, A.D., Hubbard, N.E., et al. (2018). Commonly Occurring Cell Subsets in High-Grade Serous Ovarian Tumors Identified by Single-Cell Mass Cytometry. Cell reports 22, 1875–1888.

Gutierrez-Lopez, M.D., Gilsanz, A., Yanez-Mo, M., Ovalle, S., Lafuente, E.M., Dominguez, C., Monk, P.N., Gonzalez-Alvaro, I., Sanchez-Madrid, F., and Cabanas, C. (2011). The sheddase activity of ADAM17/TACE is regulated by the tetraspanin CD9. Cellular and molecular life sciences : CMLS 68, 3275–3292.

Hamieh, M., Dobrin, A., Cabriolu, A., van der Stegen, S.J.C., Giavridis, T., Mansilla-Soto, J., Eyquem, J., Zhao, Z., Whitlock, B.M., Miele, M.M., et al. (2019). CAR T cell trogocytosis and cooperative killing regulate tumour antigen escape. Nature 568, 112–116.

Han, G., Chen, S.Y., Gonzalez, V.D., Zunder, E.R., Fantl, W.J., and Nolan, G.P. (2017). Atomic mass tag of bismuth-209 for increasing the immunoassay multiplexing capacity of mass cytometry. Cytometry A 91, 1150–1163.

Hanna, J., Goldman-Wohl, D., Hamani, Y., Avraham, I., Greenfield, C., Natanson-Yaron, S., Prus, D., Cohen-Daniel, L., Arnon, T.I., Manaster, I., et al. (2006). Decidual NK cells regulate key developmental processes at the human fetal-maternal interface. Nature medicine 12, 1065–1074.

Hanna, J., and Mandelboim, O. (2007). When killers become helpers. Trends in immunology 28, 201–206.

Hermanson, D.L., Bendzick, L., Pribyl, L., McCullar, V., Vogel, R.I., Miller, J.S., Geller, M.A., and Kaufman, D.S. (2016). Induced Pluripotent Stem Cell-Derived Natural Killer Cells for Treatment of Ovarian Cancer. Stem Cells 34, 93–101.

Horowitz, A., Strauss-Albee, D.M., Leipold, M., Kubo, J., Nemat-Gorgani, N., Dogan, O.C., Dekker, C.L., Mackey, S., Maecker, H., Swan, G.E., et al. (2013). Genetic and environmental determinants of human NK cell diversity revealed by mass cytometry. Science translational medicine 5, 208ra145.

Hotson, A.N., Gopinath, S., Nicolau, M., Khasanova, A., Finck, R., Monack, D., and Nolan, G.P. (2016). Coordinate actions of innate immune responses oppose those of the adaptive immune system during Salmonella infection of mice. Science signaling 9, ra4.

Huergo-Zapico, L., Parodi, M., Cantoni, C., Lavarello, C., Fernandez-Martinez, J.L., Petretto, A., DeAndres-Galiana, E.J., Balsamo, M., Lopez-Soto, A., Pietra, G., et al. (2018). NK-cell Editing Mediates Epithelial-to-Mesenchymal Transition via Phenotypic and Proteomic Changes in Melanoma Cell Lines. Cancer Res 78, 3913–3925.

Ideker, T., and Krogan, N.J. (2012). Differential network biology. Molecular systems biology 8, 565.

Ito, M., Nishizawa, H., Tsutsumi, M., Kato, A., Sakabe, Y., Noda, Y., Ohwaki, A., Miyazaki, J., Kato, T., Shiogama, K., et al. (2018). Potential role for nectin-4 in the pathogenesis of pre-eclampsia: a molecular genetic study. BMC medical genetics 19, 166.

Jabrane-Ferrat, N. (2019). Features of Human Decidual NK Cells in Healthy Pregnancy and During Viral Infection. Frontiers in immunology 10, 1397.

Joly, E., and Hudrisier, D. (2003). What is trogocytosis and what is its purpose? Nat Immunol 4, 815.

Kamiya, T., Seow, S.V., Wong, D., Robinson, M., and Campana, D. (2019). Blocking expression of inhibitory receptor NKG2A overcomes tumor resistance to NK cells. J Clin Invest 130, 2094–2106.

Kandalaft, L.E., Odunsi, K., and Coukos, G. (2019). Immunotherapy in Ovarian Cancer: Are We There Yet? J Clin Oncol 37, 2460–2471.

Kim, C.H., Butcher, E.C., and Johnston, B. (2002). Distinct subsets of human Valpha24-invariant NKT cells: cytokine responses and chemokine receptor expression. Trends in immunology 23, 516–519.

Koopman, L.A., Kopcow, H.D., Rybalov, B., Boyson, J.E., Orange, J.S., Schatz, F., Masch, R., Lockwood, C.J., Schachter, A.D., Park, P.J., and Strominger, J.L. (2003). Human decidual natural killer cells are a unique NK cell subset with immunomodulatory potential. J Exp Med 198, 1201–1212.

Kopcow, H.D., Allan, D.S., Chen, X., Rybalov, B., Andzelm, M.M., Ge, B., and Strominger, J.L. (2005). Human decidual NK cells form immature activating synapses and are not cytotoxic. Proc Natl Acad Sci U S A 102, 15563–15568.

Koreck, A., Suranyi, A., Szony, B.J., Farkas, A., Bata-Csorgo, Z., Kemeny, L., and Dobozy, A. (2002). CD3+CD56+ NK T cells are significantly decreased in the peripheral blood of patients with psoriasis. Clin Exp Immunol 127, 176–182.

Kotecha, N., Krutzik, P.O., and Irish, J.M. (2010). Web-based analysis and publication of flow cytometry experiments. Current protocols in cytometry / editorial board, J. Paul Robinson, managing editor … [et al.] Chapter 10, Unit10 17.

Krenning, L., van den Berg, J., and Medema, R.H. (2019). Life or Death after a Break: What Determines the Choice? Mol Cell.

Labani-Motlagh, A., Israelsson, P., Ottander, U., Lundin, E., Nagaev, I., Nagaeva, O., Dehlin, E., Baranov, V., and Mincheva-Nilsson, L. (2016). Differential expression of ligands for NKG2D and DNAM-1 receptors by epithelial ovarian cancer-derived exosomes and its influence on NK cell cytotoxicity. Tumour Biol 37, 5455–5466.

Lanier, L.L. (2015). NKG2D Receptor and Its Ligands in Host Defense. Cancer immunology research 3, 575–582.

Lee, L., and Matulonis, U. (2019). Immunotherapy and radiation combinatorial trials in gynecologic cancer: A potential synergy? Gynecol Oncol.

Li, Y., Hermanson, D.L., Moriarity, B.S., and Kaufman, D.S. (2018). Human iPSC-Derived Natural Killer Cells Engineered with Chimeric Antigen Receptors Enhance Anti-tumor Activity. Cell Stem Cell 23, 181–192.e185.

Lopez-Soto, A., Gonzalez, S., Smyth, M.J., and Galluzzi, L. (2017). Control of Metastasis by NK Cells. Cancer Cell 32, 135–154.

Lord, C.J., and Ashworth, A. (2017). PARP inhibitors: Synthetic lethality in the clinic. Science 355, 1152–1158.

Lorenzo-Herrero, S., Lopez-Soto, A., Sordo-Bahamonde, C., Gonzalez-Rodriguez, A.P., Vitale, M., and Gonzalez, S. (2018). NK Cell-Based Immunotherapy in Cancer Metastasis. Cancers 11.

Lorenzo-Herrero, S., Sordo-Bahamonde, C., Gonzalez, S., and Lopez-Soto, A. (2020). Evaluation of NK cell cytotoxic activity against malignant cells by the calcein assay. Methods Enzymol 631, 483–495.

Martinet, L., and Smyth, M.J. (2015). Balancing natural killer cell activation through paired receptors. Nat Rev Immunol 15, 243–254.

Matulonis, U.A. (2018). Management of newly diagnosed or recurrent ovarian cancer. Clin Adv Hematol Oncol 16, 426–437.

Matulonis, U.A., Sood, A.K., Fallowfield, L., Howitt, B.E., Sehouli, J., and Karlan, B.Y. (2016). Ovarian cancer. Nat Rev Dis Primers 2, 16061.

McCann, F.E., Eissmann, P., Onfelt, B., Leung, R., and Davis, D.M. (2007). The activating NKG2D ligand MHC class I-related chain A transfers from target cells to NK cells in a manner that allows functional consequences. J Immunol 178, 3418–3426.

Miyake, K., Shiozawa, N., Nagao, T., Yoshikawa, S., Yamanishi, Y., and Karasuyama, H. (2017). Trogocytosis of peptide-MHC class II complexes from dendritic cells confers antigen-presenting ability on basophils. Proc Natl Acad Sci U S A 114, 1111–1116.

Morvan, M.G., and Lanier, L.L. (2016). NK cells and cancer: you can teach innate cells new tricks. Nat Rev Cancer 16, 7–19.

Neri, S., Mariani, E., Meneghetti, A., Cattini, L., and Facchini, A. (2001). Calcein-acetyoxymethyl cytotoxicity assay: standardization of a method allowing additional analyses on recovered effector cells and supernatants. Clinical and diagnostic laboratory immunology 8, 1131–1135.

Okumura, G., Iguchi-Manaka, A., Murata, R., Yamashita-Kanemaru, Y., Shibuya, A., and Shibuya, K. (2020). Tumor-derived soluble CD155 inhibits DNAM-1-mediated antitumor activity of natural killer cells. J Exp Med 217.

Orr, M.T., and Lanier, L.L. (2010). Natural killer cell education and tolerance. Cell 142, 847–856.

Quah, B.J., Barlow, V.P., McPhun, V., Matthaei, K.I., Hulett, M.D., and Parish, C.R. (2008). Bystander B cells rapidly acquire antigen receptors from activated B cells by membrane transfer. Proc Natl Acad Sci U S A 105, 4259–4264.

Ramadoss, N.S., Zhao, N.Q., Richardson, B.A., Grant, P.M., Kim, P.S., and Blish, C.A. (2019). Enhancing natural killer cell function with gp41-targeting bispecific antibodies to combat HIV infection. bioRxiv, 760280.

Raulet, D.H., Gasser, S., Gowen, B.G., Deng, W., and Jung, H. (2013). Regulation of ligands for the NKG2D activating receptor. Annu Rev Immunol 31, 413–441.

Reches, A., Ophir, Y., Stein, N., Kol, I., Isaacson, B., Charpak Amikam, Y., Elnekave, A., Tsukerman, P., Kucan Brlic, P., Lenac, T., et al. (2020). Nectin4 is a novel TIGIT ligand which combines checkpoint inhibition and tumor specificity. J Immunother Cancer 8.

Reed, J., and Wetzel, S.A. (2019). Trogocytosis-Mediated Intracellular Signaling in CD4(+) T Cells Drives TH2-Associated Effector Cytokine Production and Differentiation. J Immunol 202, 2873–2887.

Reyes, R., Cardenes, B., Machado-Pineda, Y., and Cabanas, C. (2018). Tetraspanin CD9: A Key Regulator of Cell Adhesion in the Immune System. Frontiers in immunology 9, 863.

Rezvani, K. (2019). Adoptive cell therapy using engineered natural killer cells. Bone Marrow Transplant 54, 785–788.

Rezvani, K., Rouce, R., Liu, E., and Shpall, E. (2017). Engineering Natural Killer Cells for Cancer Immunotherapy. Mol Ther 25, 1769–1781.

Rodriguez, G.M., and Galpin, K.J.C. (2018). The Tumor Microenvironment of Epithelial Ovarian Cancer and Its Influence on Response to Immunotherapy. 10.

Samusik, N., Good, Z., Spitzer, M.H., Davis, K.L., and Nolan, G.P. (2016). Automated mapping of phenotype space with single-cell data. Nat Methods.

Sanchez-Correa, B., Valhondo, I., Hassouneh, F., Lopez-Sejas, N., Pera, A., Bergua, J.M., Arcos, M.J., Banas, H., Casas-Aviles, I., Duran, E., et al. (2019). DNAM-1 and the TIGIT/PVRIG/TACTILE Axis: Novel Immune Checkpoints for Natural Killer Cell-Based Cancer Immunotherapy. Cancers 11.

Shabrish, S., Gupta, M., and Madkaikar, M. (2016). A Modified NK Cell Degranulation Assay Applicable for Routine Evaluation of NK Cell Function. Journal of immunology research 2016, 3769590.

Siebert, J.C., Inokuma, M., Waid, D.M., Pennock, N.D., Vaitaitis, G.M., Disis, M.L., Dunne, J.F., Wagner, D.H., Jr., and Maecker, H.T. (2008). An analytical workflow for investigating cytokine profiles. Cytometry A 73, 289–298.

Singh, N., McCluggage, W.G., and Gilks, C.B. (2017). High-grade serous carcinoma of tubo-ovarian origin: recent developments. Histopathology 71, 339–356.

Spitzer, M.H., Carmi, Y., Reticker-Flynn, N.E., Kwek, S.S., Madhireddy, D., Martins, M.M., Gherardini, P.F., Prestwood, T.R., Chabon, J., Bendall, S.C., et al. (2017). Systemic Immunity Is Required for Effective Cancer Immunotherapy. Cell.

Suck, G., Linn, Y.C., and Tonn, T. (2016). Natural Killer Cells for Therapy of Leukemia. Transfusion medicine and hemotherapy : offizielles Organ der Deutschen Gesellschaft fur Transfusionsmedizin und Immunhamatologie 43, 89–95.

Tabiasco, J., Espinosa, E., Hudrisier, D., Joly, E., Fournie, J.J., and Vercellone, A. (2002). Active trans-synaptic capture of membrane fragments by natural killer cells. Eur J Immunol 32, 1502–1508.

Tilburgs, T., Evans, J.H., Crespo, A.C., and Strominger, J.L. (2015). The HLA-G cycle provides for both NK tolerance and immunity at the maternal-fetal interface. Proc Natl Acad Sci U S A 112, 13312–13317.

Tremblay-McLean, A., Coenraads, S., Kiani, Z., Dupuy, F.P., and Bernard, N.F. (2019). Expression of ligands for activating natural killer cell receptors on cell lines commonly used to assess natural killer cell function. BMC immunology 20, 8.

Uppendahl, L.D., Dahl, C.M., Miller, J.S., Felices, M., and Geller, M.A. (2017). Natural Killer Cell-Based Immunotherapy in Gynecologic Malignancy: A Review. Frontiers in immunology 8, 1825.

Venkitaraman, A.R. (2009). Linking the cellular functions of BRCA genes to cancer pathogenesis and treatment. Annual review of pathology 4, 461–487.

Vento-Tormo, R., Efremova, M., Botting, R.A., Turco, M.Y., Vento-Tormo, M., Meyer, K.B., Park, J.E., Stephenson, E., Polanski, K., Goncalves, A., et al. (2018). Single-cell reconstruction of the early maternal-fetal interface in humans. Nature 563, 347–353.

Vivier, E., Raulet, D.H., Moretta, A., Caligiuri, M.A., Zitvogel, L., Lanier, L.L., Yokoyama, W.M., and Ugolini, S. (2011). Innate or adaptive immunity? The example of natural killer cells. Science 331, 44–49.

Vyas, J.M., Van der Veen, A.G., and Ploegh, H.L. (2008). The known unknowns of antigen processing and presentation. Nat Rev Immunol 8, 607–618.

Wilk, A.J., and Blish, C.A. (2018). Diversification of human NK cells: Lessons from deep profiling. J Leukoc Biol 103, 629–641.

Wroblewski, E.E., Parham, P., and Guethlein, L.A. (2019). Two to Tango: Co-evolution of Hominid Natural Killer Cell Receptors and MHC. Frontiers in immunology 10, 177.

Yoneda, J., Kuniyasu, H., Crispens, M.A., Price, J.E., Bucana, C.D., and Fidler, I.J. (1998). Expression of angiogenesis-related genes and progression of human ovarian carcinomas in nude mice. J Natl Cancer Inst 90, 447–454.

Yurtsever, A., Haydaroglu, A., Biray Avci, C., Gunduz, C., Oktar, N., Dalbasti, T., Caglar, H.O., Attar, R., and Kitapcioglu, G. (2013). Assessment of genetic markers and glioblastoma stem-like cells in activation of dendritic cells. Human cell 26, 105–113.

Zhang, A.W., McPherson, A., Milne, K., Kroeger, D.R., Hamilton, P.T., Miranda, A., Funnell, T., Little, N., de Souza, C.P.E., Laan, S., et al. (2018). Interfaces of Malignant and Immunologic Clonal Dynamics in Ovarian Cancer. Cell 173, 1755–1769.e1722.

Zhang, Y., and Weinberg, R.A. (2018). Epithelial-to-mesenchymal transition in cancer: complexity and opportunities. Frontiers of medicine 12, 361–373.

Zoller, T., Wittenbrink, M., Hoffmeister, M., and Steinle, A. (2018). Cutting an NKG2D Ligand Short: Cellular Processing of the Peculiar Human NKG2D Ligand ULBP4. Frontiers in immunology 9, 620.

