## Supplemental material for "High-Grade Serous Ovarian Tumor Cells Modulate NK Cell Function to Create an Immune-Tolerant Microenvironment"

### Supplemental Information

### Supplemental Figures

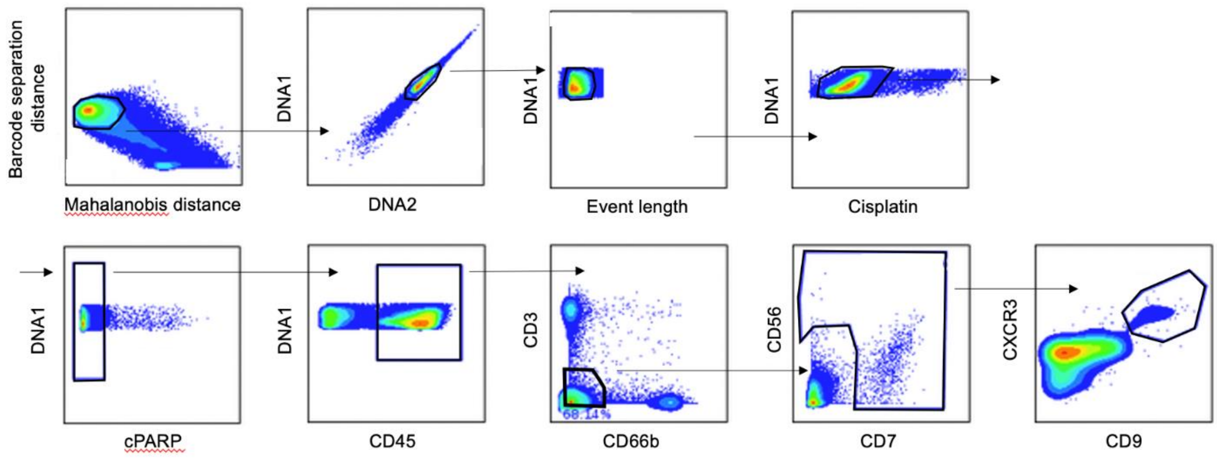


**Figure S1. Manual gating scheme for dl-NK cells.** dl-NK cells gated from the CD45+ immune cell infiltrate shown here from a representative HGSC sample.

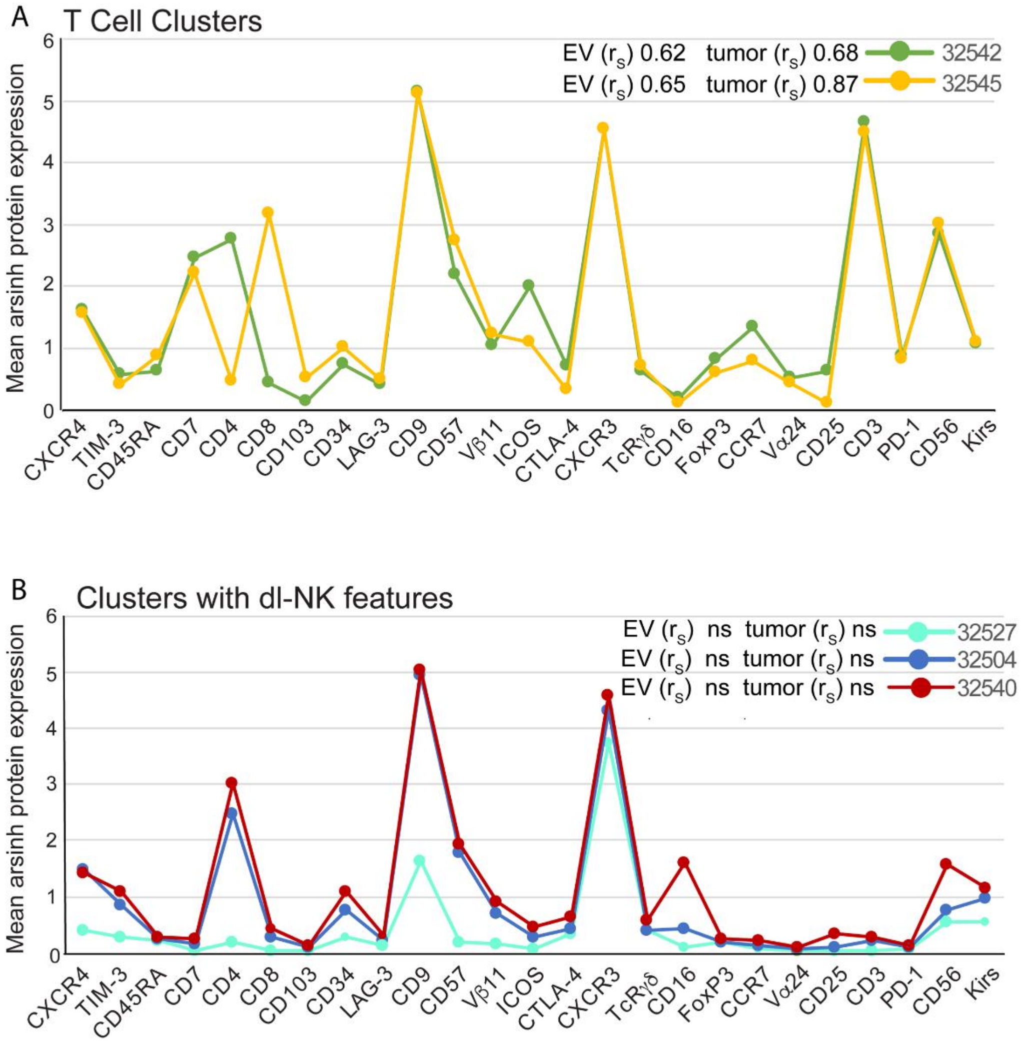


**Figure S2. Cell clusters with dl-like and T cell phenotypes.** T cell clusters correlated with total and EV tumor cells. **(B)** Three clusters with dl-NK cell features that did not correlate with total tumor or EV tumor cells but were found in all tumors. Clusters 32504 and 32540 were present with a combined frequency range of 14% - 86% of the immune cell infiltrate (CD45+ CD66-).

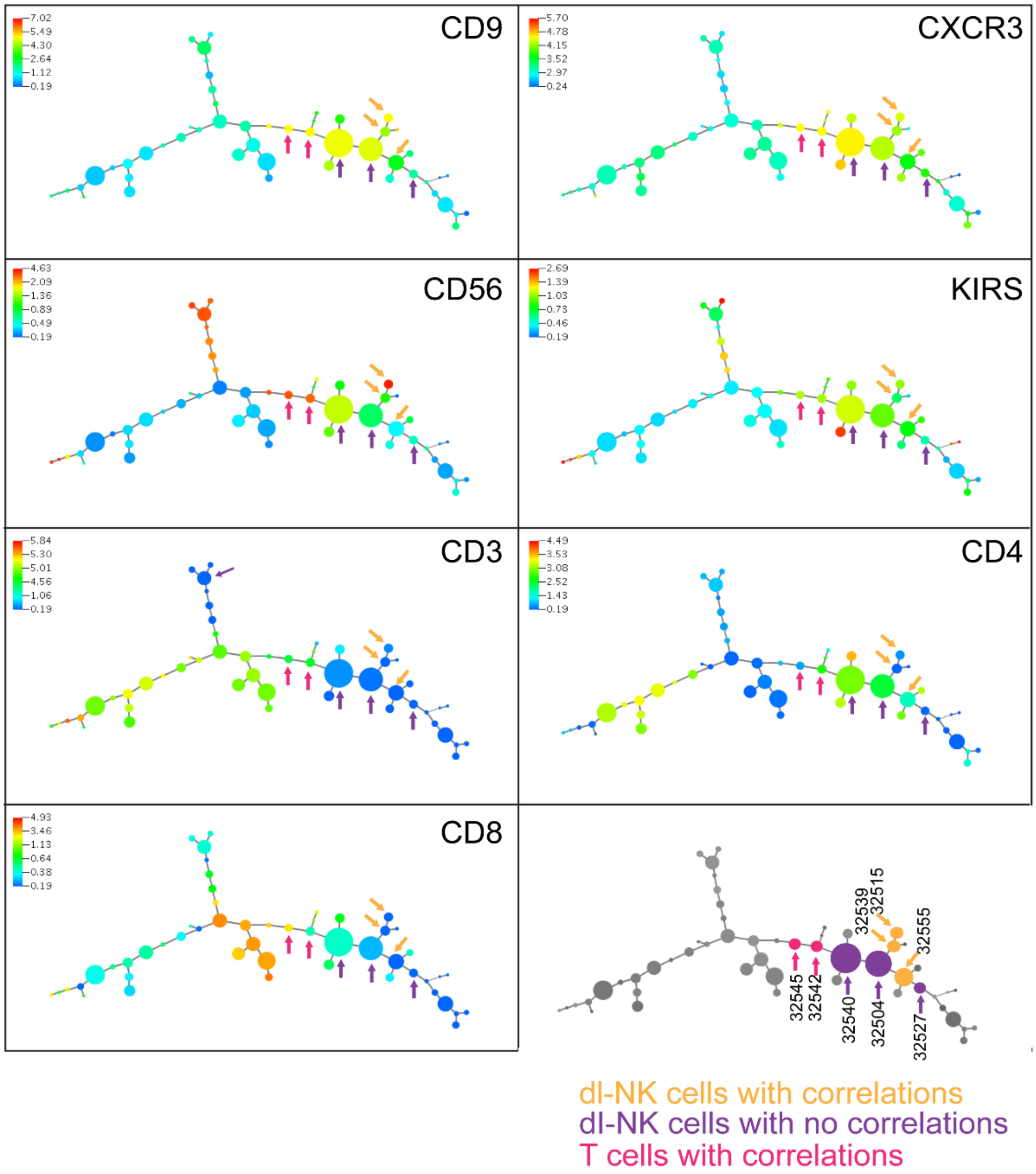


**Figure S3. Composite minimum spanning trees (MST) of NK and T cell X-shift clusters from HGSC tumor infiltrate**. The 56 X-shift clusters generated by the X-shift algorithm were visualized along an MST such that neighboring clusters had similar marker co-expression patterns (Samusik et al., 2016). The MST in the bottom right is a key and depicts the seven dl-NK cell clusters we identified with and without correlations with tumor cell abundance and total EV cell abundance (Figure 1B and Figure S2).

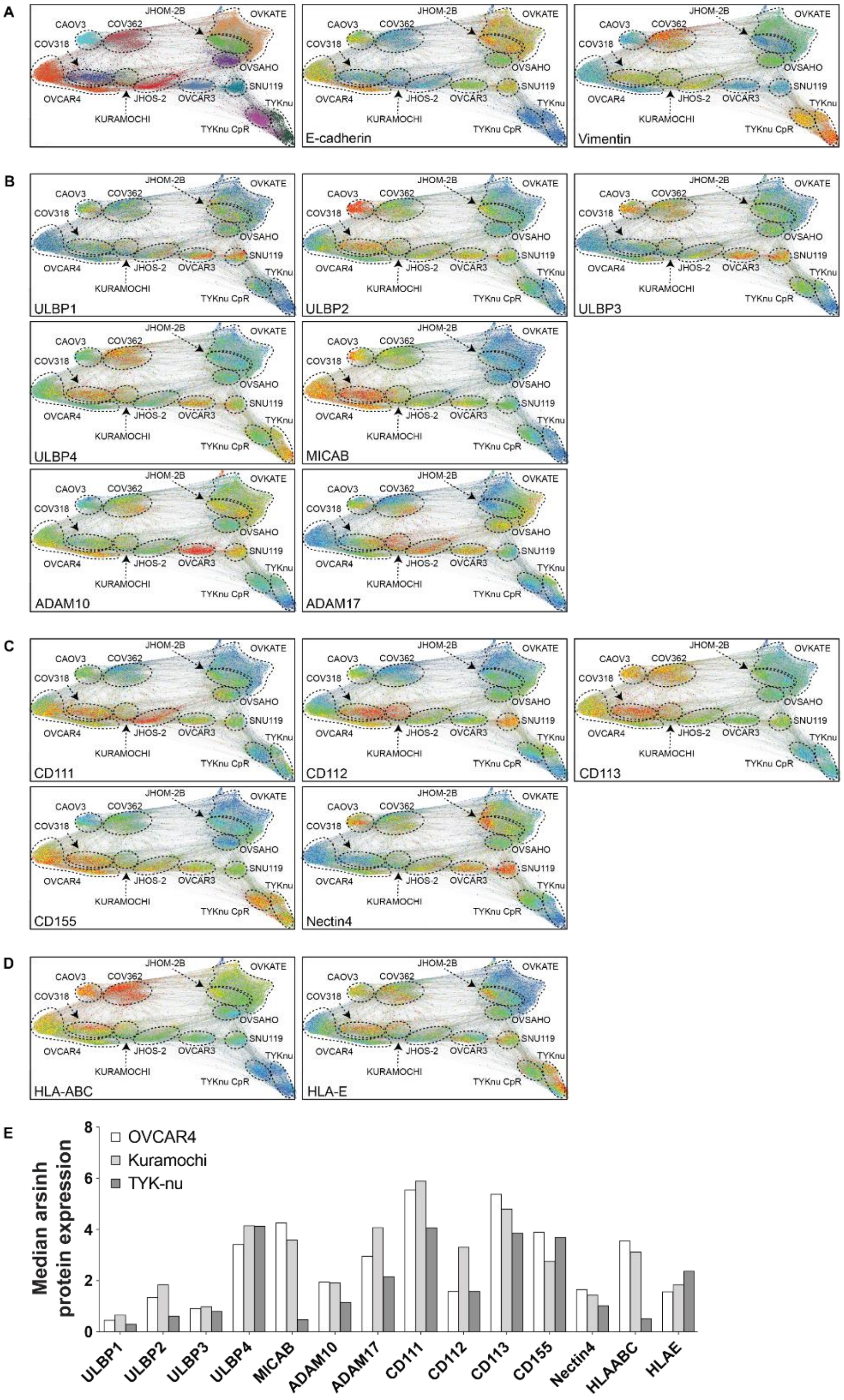


**Figure S4. Expression patterns of NK receptor ligands in HGSC cell lines.** Single cell force directed layouts (FDLs) of 10,000 cells sampled from X-shift clusters of HGSC cell lines colored for: **(A)** Cell line (far left panel), expression levels of E-cadherin and vimentin. **(B)** NKG2D ligands and ADAM10 and 17. **(C)** Nectin family ligands. **(D)** HLA-ABC and HLA-E that bind the inhibitory KIRs and NKG2A/CD94 receptors, respectively. **(E)** Bar chart shows expression of each protein in the OVCAR4, Kuramochi and TYK-nu HGSC cell lines.

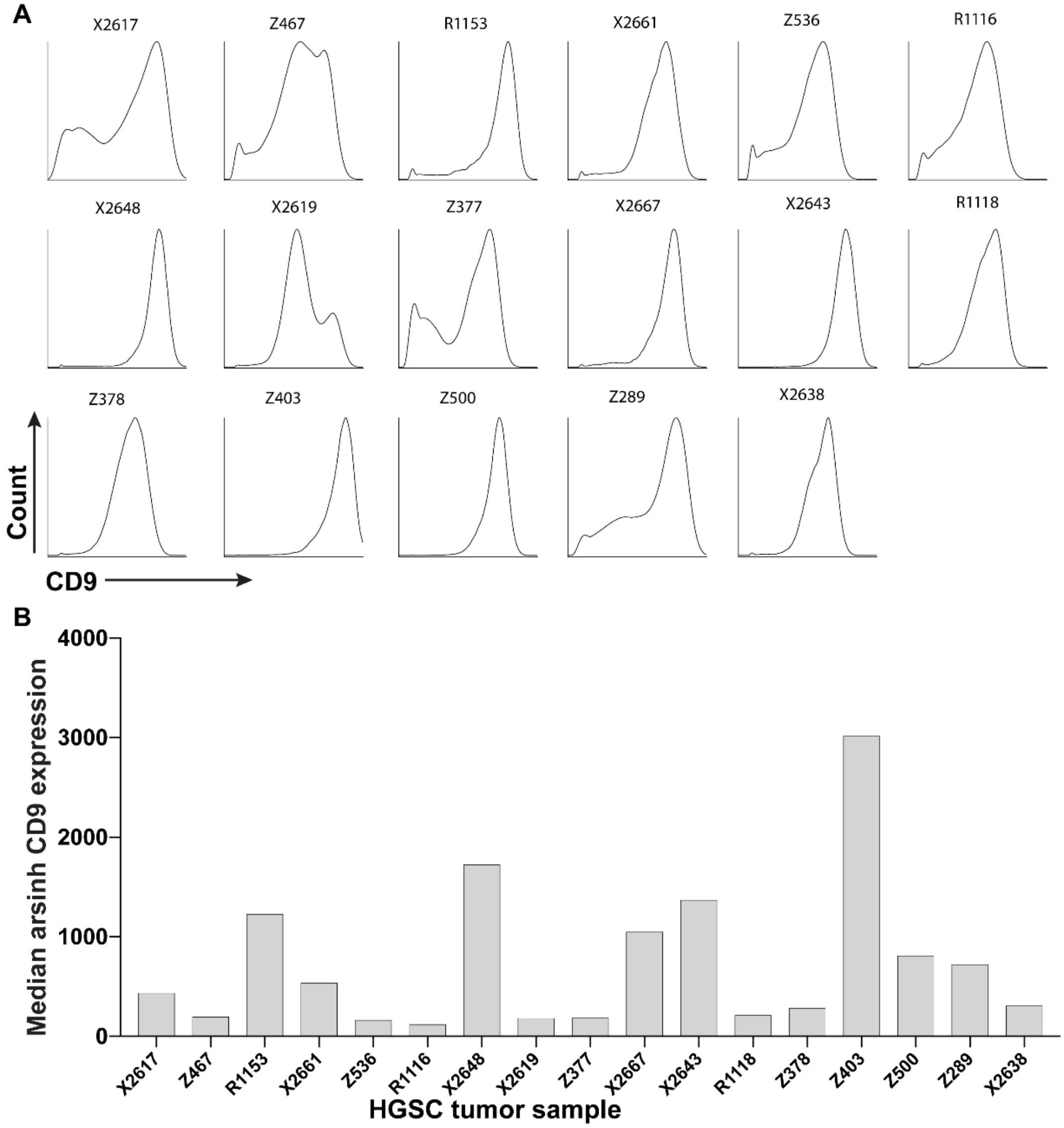


**Figure S5. CD9 expression levels in primary tumors.** Tumor cells from each newly diagnosed tumor (Table S1) were gated as CD45- FAP- CD31- as descried previously(Gonzalez et al., 2018). Variable but high levels of CD9 were expressed on HGSC tumor cells across all samples. (A) frequencies of HGSC tumor cells expressing CD9. (B) Median expression levels of CD9 across tumors.

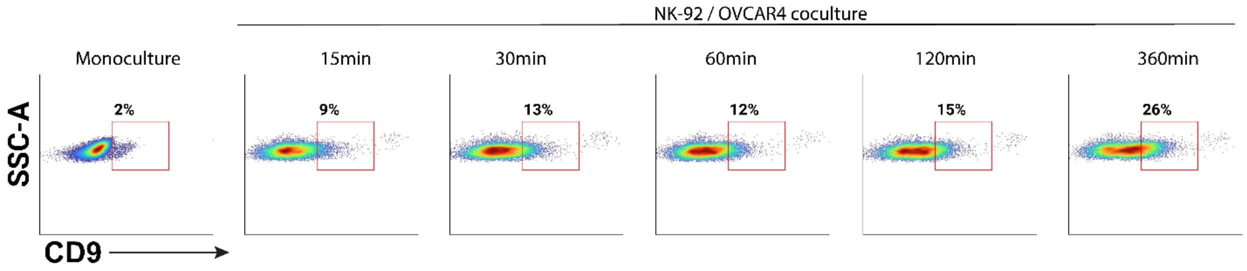


**Figure S6. Time course for CD9 expression on NK-92 cells.** OVCAR4 and NK-92 cells were cocultured at a 1:1 ratio. At the times indicated, CD9 expression levels were measured on the NK-92 cells by flow cytometry. An antibody against CD45 was used to gate the NK-92 cell population from the coculture.

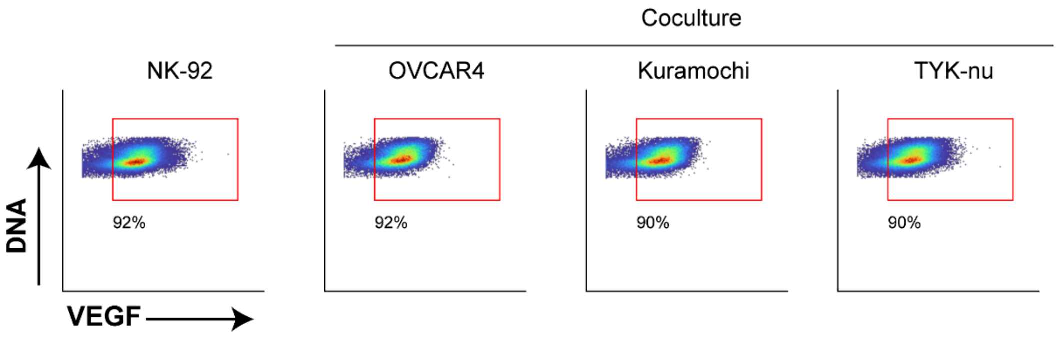


| Expression of VEGF (raw median values) | | | |
|---|---|---|---|
| NK92 monoculture | NK92: OVCAR4 | NK92: Kuramochi | NK92: TYKnu |
| 30.26 | 34.96 | 30.71 | 29.19 |

**Figure S7. VEGF is consitutively expressed by the NK-92 cell line.** High frequencies of VEGF-expressing NK-92 cells grown in monoculture with no significant changes in exresssion levels after coculture with HGSC cell lines. Representative experiment shown.

### Supplemental Tables

| Case no | Patient ID | Age | Localization | Histological type | Stage | Radicality | Grade | Morphology |
|---|---|---|---|---|---|---|---|---|
| 1 | X2617 | 63 | ovary | ovarian carcinoma - serous type | III C | R2 | G3 | serous cystadenocarcinoma, NOS |
| 2 | X2619 | 57 | ovary | ovarian carcinoma - serous type | IIC | R0 | G2 | serous cystadenocarcinoma, NOS |
| 3 | Z289 | 79 | ovary | ovarian carcinoma - serous papillary type | III C | RX | G3 | papillary serous cystadenocarcinoma |
| 4 | X2648* | 67 | peritoneum | ovarian carcinoma - serous papillary type | III C | R1 | G3 | serous cystadenocarcinoma, NOS (C56.9) |
| 5 | Z378 | 71 | ovary | ovarian carcinoma - serous papillary type | III C | R0 | G3 | serous cystadenocarcinoma, NOS (C56.9) |
| 6 | X2643 | 53 | peritoneum | ovarian carcinoma - serous papillary type | III B | R0 | G3 | serous cystadenocarcinoma, NOS (C56.9) |
| 7 | Z403 | 82 | ovary | ovarian carcinoma | III C | RX | G3 | serous surface papillary carcinoma (C56.9) |
| 8 | X2638 | 72 | ovary | adenocarcinoma | IV | R2 | G3 | adenocarcinoma, NOS |
| 9 | Z377† | 56 | ovary | ovarian carcinoma - serous papillary type | IV | R0 | G3 | serous surface papillary carcinoma |
| 10 | Z467 | 48 | omentum | ovarian carcinoma - serous papillary type | III C | R0 | G3 | serous cystadenocarcinoma, NOS (C56.9) |
| 11 | R1116 | 67 | ovary | ovarian carcinoma - serous papillary type | III C | R1 | G3 | serous surface papillary carcinoma (C56.9) |
| 12 | R1118 | 84 | ovary | ovarian carcinoma - serous type | III A | RX | G3 | serous cystadenocarcinoma, NOS (C56.9) |
| 13 | Z500 | 58 | ovary | ovarian carcinoma - serous papillary type | III C | R0 | G3 | serous surface papillary carcinoma (C56.9) |
| 14 | X2661* | 74 | peritoneum | ovarian carcinoma - serous papillary type | III C | R2 | G3 | serous surface papillary carcinoma (C56.9) |
| 15 | R1153 | 88 | ovary | ovarian carcinoma - serous papillary type | III C | R2 | G3 | papillary serous cystadenocarcinoma (C56.9) |
| 16 | Z536 | 82 | omentum | ovarian carcinoma - serous papillary type | III C | RX | G3 | serous cystadenocarcinoma, NOS (C56.9) |
| 17 | X2667 | 69 | ovary | ovarian carcinoma - serous papillary type | III C | R0 | G3 | serous surface papillary carcinoma (C56.9) |
| 18 | Z573 |  | omentum | ovarian carcinoma - serous papillary type | III C | RX | G3 | serous surface papillary carcinoma (C56.9) |
| 19 | R1181 | 84 | ovary | ovarian carcinoma - serous papillary type | III C | R0 | G3 | serous surface papillary carcinoma (C56.9) |
| 20 | R1283 | 74 | omentum | ovarian carcinoma - serous papillary type | n/a | R1 | n/a | papillary serous cystadenocarcinoma (C56.9) |

**Table S1. Clinical characteristics of patient samples.** Newly diagnosed chemo-naïve tumors were all diagnosed as malignant neoplasms of the ovary. All patients underwent surgical de-bulking followed by a platinum-based chemotherapeutic regimen. Samples 1 to 17 were analyzed in our prior study that studied the tumor cells(Gonzalez et al., 2018). For this study, the same samples 1 to 17 were analyzed with a CyTOF panel that characterized intra-tumoral T and NK cells. Green highlight indicates samples analyzed with a CyTOF panel designed to interrogate NK receptor ligands. NOS – not otherwise specified.

| Case # | Patient ID | Symbol | Chrom | Position | rs ID | REF | ALT | Filter | Consequence | COSMIC ID | Clinvar status | AA or cDNA | SIFT prediction | Variant read fraction |
|---|---|---|---|---|---|---|---|---|---|---|---|---|---|---|
| **1.** | X2648 | TP53 | chr17 | 7577593 | | TAC | T | PASS | frameshift | COSM111635 | NA | p.C229fs*10 | NA | 0.333 |
| **2.** | R1181 | TP53 | chr17 | 7577121 | rs121913343 | G | A | PASS | missense | COSM10659 | pathogenic | p.R273C | deleterious | 0.853 |
| **3.** | X2661 | TP53 | chr17 | 7579358 | rs11540654 | C | G | PASS | missense | COSM10716 | NA | p.R110L | deleterious | 0.674 |
| **4.** | R1283 | TP53 | chr17 | 7577535 | | CCTCCGGTTCATGCCGCCCATGCAGGAACTGTTACA | C | PASS | deletion | | NA | c.712_746del | | 0.135 |
| **5.** | Z536 | TP53 | chr17 | 7577538 | rs11540652 | C | T | PASS | missense | COSM10662 | pathogenic | p.R248Q | deleterious | 0.816 |
| **6.** | Z403 | TP53 | chr17 | 7578394 | | T | A | PASS | missense | COSM10889 | NA | p.H179R | deleterious | 0.741 |
| **6.** | Z403 | **BRCA2** | chr13 | 32936731 | | G | A | PASS | stop gained | NA | pathogenic | p.W2626X | | 0.609 |
| **7.** | Z377 | TP53 | chr17 | 7578403 | | C | T | PASS | missense | COSM10645 | NA | p.C176F | deleterious | 0.255 |
| **7.** | Z377 | **BRCA2** | chr13 | 32900672 | | G | T | PASS | stop gained | NA | pathogenic | p.G185X | | 0.231 |
| **8.** | Z378 | TP53 | chr17 | 7578290 | | C | T | PASS | splice acceptor | COSM127200 | NA | c.560-1G>T | NA | 0.622 |
| **9.** | Z573 | TP53 | chr17 | 7578203 | rs730882025 | C | T | PASS | missense | COSM3388195 | pathogenic | p.V216M | deleterious | 0.662 |
| **9.** | Z573 | **BRCA1** | chr17 | 41243713 | rs1060505047 | GC | G | PASS | frameshift | | pathogenic | p.A1279Hfs | NA | 0.855 |
| **10.** | R1118 | TP53 | chr17 | 7577548 | rs28934575 | C | A | PASS | missense | COSM10957 | pathogenic | p.G245R | deleterious | 0.96 |
| **11.** | X2643 | TP53 | chr17 | 7577082 | | C | T | PASS | missense | COSM10726 | NA | p.E286K | deleterious | 0.659 |
| **12.** | X2638 | TP53 | chr17 | 7577520 | | A | C | PASS | missense | COSM1230106 | NA | p.I254S | deleterious | 0.826 |
| **13.** | Z467 | TP53 | chr17 | 7578190 | rs121912666 | T | C | PASS | missense | COSM10758 | pathogenic | p.Y220C | deleterious | 0.14 |
| **15.** | Z500 | TP53 | chr17 | 7578190 | rs121912666 | T | C | PASS | missense | COSM10758 | pathogenic | p.Y220C | deleterious | 0.785 |
| **18.** | X2617 | TP53 | chr17 | 7579349 | | A | C | PASS | missense | COSM10717 | | p.F113C | deleterious | 0.13 |
| **19.** | X2619 | TP53 | chr17 | 7577539 | rs121912651 | G | A | PASS | missense | COSM10656 | pathogenic | p.R248W | deleterious | 0.818 |
| **20.** | Z289 | TP53 | cr17 | 7577539 | rs121912651 | G | A | PASS | missense | COSM10656 | pathogenic | p.R248W | deleterious | 0.286 |

**Table S2. Pathogenic or predicted deleterious variants of TP53, BRCA1 and BRCA2.** Sequencing was performed as described in Materials and Methods. Samples were characterized as described above in legend to Table S1. Material unavailable for sequencing samples 14, 16 and 17.

| Isotope | Antigen | Clone | Phospho site | Vendor | Positive control | Negative control |
|---|---|---|---|---|---|---|
| Y 89 | CD45 | H130 | | Fluidigm | U937 | OVCAR3 |
| In 113 | FAP | F11-24 | | eBioscience | WI-38 | Ramos |
| In 115 | **CXCR4** | 12G5 | | Biolegend | NIH3T3 | PBMC - CD8+ T cells |
| Ce 140 | CD31 | WM59 | | Biolegend | HUVEC | U937 |
| Pr 141 | **TIM-3** | 344823 | | R&D systems | PBMC - CD8+ T cells | PBMC - CD14+ Monocytes |
| Nd 142 | pMAPKAPK2 | 27B7 | pT334 | CST | PBMC - CD14+ Monocytes LPS 30min | PBMC - CD14+ Monocytes dd H2O |
| Nd 143 | **CD45RA** | HI100 | | Fluidigm | PBMC - CD8+ CD45RO- | PBMC - CD8+ CD45RO+ |
| Nd 144 | **CD7** | M-T701 | | BD Pharmingen | PBMC - CD4+ T cells | PBMC - CD14+ Monocytes |
| Nd 145 | **CD4** | RPA-T4 | | Fluidigm | PBMC - CD3+CD8- cells | PBMC - CD3+CD8+ cells |
| Nd 146 | **CD8a** | RPA-T8 | | Fluidigm | PBMC - CD3+CD4- cells | PBMC - CD3+CD4+ cells |
| Sm 147 | **CD103** | Ber-ACT8 | | BD | PBMC- CD3+ PHA 72hr | PBMC- CD3+ PBS |
| Nd 148 | **CD34** | 583 | | Fluidigm | OVCAR3, K562 | U937 |
| Sm 149 | pNFkB | K10-895.12.50 | pS529 | BD | Jurkat_TNFa (10ng/ml, 15min) | Jurkat_PBS |
| Nd 150 | **LAG-3 (CD223)** | 874501 | | Fluidigm | PBMC - CD8+ PMA/Iono 24hr | PBMC - CD8+ Ethanol (0.2%) |
| Eu 151 | Ki67 (total) | B56 | | BD | OVCAR4 | OVKATE |
| Sm 152 | CD66b | G10F5 | | Biolegend | Whole Blood -CD45low | Whole Blood - CD3+ T cells |
| Eu 153 | pSTAT1 | 58D6 | pY701 | Fluidigm | PBMC - CD14+ Monocytes IFNg 15min | PBMC - CD14+ Monocytes PBS |
| Sm 154 | pSTAT3 | 4/P-STAT3 | pY705 | BD | Jurkat_Pervanadate (125uM, 15min) | Jurkat_DMSO (0.0625%) |
| Gd 155 | IkB-alpha Total (N-term antigen) | L35A5 | | CST | PBMC - CD14+ Monocytes LPS 30min | PBMC - CD14+ Monocytes ddH2O |
| Gd 156 | **CD9** | M-L13 | | BD | Whole Blood - CD61+ Platelete | Whole Blood - CD3+ T cells |
| Gd 157 | **CD57** | HCD57 | | Biolegend | PBMC - CD8+ T cells: CD45RA+CCR7- | PBMC - CD19+ B cells |
| Gd 158 | **Vβ11** | C21 | | Beckman coulter | PBMC - NKT cells | PBMC - CD19+ B cells |
| Tb 159 | pAkt | D9E | pS473 | CST | Kuramochi_EGF(100ng/ml, 10min) | Kuramochi_serum starved overnight |
| Gd 160 | **ICOS (CD278)** | C398.4A | | Biolegend | PBMC - CD8+ PMA/Iono 24hr | PBMC - CD8+ Ethanol (0.2%) |
| Dy 161 | **CTLA4 (CD152)** | 14D3 | | Fluidigm | PBMC - CD4+ PMA/Iono 6h, Brefeldin A +Monensin added 1h post stim | PBMC - CD4+ Ethanol (0.2%), Brefeldin A+Monensin added 1h post stim |
| Dy 162 | pSTAT5 | 47/Stat5 | pY694 | BD | Jurkat_Pervanadate (125uM, 15min) | Jurkat_DMSO (0.0625%) |
| Dy 163 | **CXCR3 (CD183)** | G025H7 | | Fluidigm | HUVEC | PBMC - CD19+ B cells and CD3+ T cells |
| Dy 164 | **TCRgd** | B1 | | Biolegend | PBMC - CD3+CD4-CD8- | PBMC - CD19+ B cells |
| Ho 165 | **CD16** | B73.1 | | eBioscience | PBMC - CD3- | PBMC - CD4+ T cells |
| Er 166 | **FoxP3** | PCH101 | | eBioscience | PBMC - CD4+ T cells | PBMC - CD8+ T cells |
| Er 167 | **CCR7** | 150503 | | R&D systems | PBMC - CD8+ CD45RA+ | PBMC - CD8+ CD45RA- |
| Er 168 | **Vα24** | C15 | | Beckman coulter | PBMC - NKT cells | PBMC - CD19+ B cells |
| Tm 169 | **CD25** | 2A3 | | Fluidigm | PBMC - CD8+ PMA/Iono 24hr | PBMC - CD8+ Ethanol (0.2%) |
| Er 170 | **CD3** | UCHT1 | | Fluidigm | PBMC - CD33-CD14- | PBMC - CD14+ Monocytes |
| Yb 171 | cPARP | F21-852 | cleaved N214 | BD | U937_Etoposide (20uM, 24hrs) | U937_DMSO (0.05%) |
| Yb 172 | prpS6 | N7-548 | pS235/pS236 | BD | OVCAR4 | U937 |
| Yb 173 | pPLCg2 | K86-689.37 | pY759 | BD | Ramos- anti-IgM 10min | Ramos - PBS |
| Yb 174 | pCREB | 87G3 | pS133 | CST | U937_PMA (400nM, 10min) | U937_DMSO (0.02%) |
| Lu 175 | **PD-1 (CD279)** | EH12.2H7 | | Fluidigm | PBMC - CD8+ PMA/Iono 72hr | PBMC - CD8+ Ethanol (0.2%) |
| Yb 176 | **CD56** | NCAM16.2 | | Fluidigm | PBMC - CD3-CD14-CD16+ | PBMC - CD4+ T cells |
| Bi 209 | anti-PE | PE001 | | Biolegend | | |
| Bi 209 | **NKB1 -PE** | DX9 | | BD Pharmingen | PBMC - CD56+ NK cells | PBMC - CD14+ Monocytes |
| Bi 209 | **CD158a -PE** | HP-3E4 | | BD Pharmingen | PBMC - CD56+ NK cells | PBMC - CD14+ Monocytes |
| Bi 209 | **CD158b -PE** | CH-L | | BD Pharmingen | PBMC - CD56+ NK cells | PBMC - CD14+ Monocytes |

**Table S3. Antibody panel designed to characterize intra-tumoral T and NK cells**. The immune infiltrate was gated from viable singlets as CD45+ CD66-. The antibodies shown in blue denote markers used for the clustering.

| Isotope | Antigen | Clone | Phospho site | Vendor | Positive control cell line | Negative control cell line |
|---|---|---|---|---|---|---|
| Y 89 | CD45 | H130 | | Fluidigm | U937 | OVCAR3 |
| In 113 | FAP | F11-24 | | eBioscience | WI-38 | Ramos |
| In 115 | **Vimentin** | D21H3 | | CST | NIH3T3 | OVCAR3 |
| Ce 140 | CD31 | WM59 | | Biolegend | HUVEC | U937 |
| Pr 141 | **CD73** | AD2 | | BD | COV362 | Ramos |
| Nd 142 | **CD61** | VI-PL2 | | BD | PBMC - CD14+ Monocytes | PBMC - CD3+ T cells |
| Nd 143 | **MUC16/CA125** | X75 (3C8/4) | | Gen Way | OVCAR3 | U937 |
| Nd 144 | HLA-ABC | W6/32 | | Fluidigm | CAVO3 | NIH3T3 |
| Nd 145 | **CD151** | 50-6 | | Biolegend | OVCAR4 | Ramos |
| Nd 146 | CD111 | R1.302 | | Biolegend | JHOS2 | OVKATE |
| Sm 147 | pH2AX | JBW301 | pS139 | Millipore | HeLa_Etoposide (10 uM, 24hrs) | HeLa_DMSO (0.05%) |
| Nd 148 | HLAE | 3D12 | | eBioscience | CAVO3 | NIH3T3 |
| Sm 149 | CD113 | N3.12.4 | | Santa Cruz Biotech | KURAMOCHI | NIH3T3 |
| Nd 150 | ULBP2/5/6 | 165903 | | R&D system | CAOV3 | TykNU |
| Eu 151 | ULBP4 | 709116 | | R&D system | TykNU | NIH3T3 |
| Sm 152 | CD155 | TX24 | | Biolegend | TykNU | NIH3T3 |
| Eu 153 | **CD49f** | MP4F10 | | R&D system | CaCo | Ramos |
| Sm 154 | MICA/B | 6D4 | | Biolegend | HeLa | TykNU |
| Gd 155 | **CD133** | AC133 | | Miltenyi | CaCo | Ramos |
| Gd 156 | **CD10** | Hl10a | | Biolegend | REH | U937 |
| Gd 157 | Nectin-4 | 337516 | | R&D system | SNU119 | NIH3T3 |
| Gd 158 | **E-Cadherin** | 67A4 | | Biolegend | OVCAR3 | U937 |
| Tb 159 | **CD90** | 5E10 | | Biolegend | OVKATE | U937 |
| Gd 160 | ULBP3 | 2F9 | | Santa Cruz Biotech | CAOV3 | OVKATE |
| Dy 161 | c-Myc | D84C12 | | CST | TykNU | MCF7 |
| Dy 162 | pSTAT5 | 47/Stat5 | pY694 | BD | Jurkat_Pervanadate (125uM, 15min) | Jurkat_DMSO (0.0625%) |
| Dy 163 | **Endoglin** | 43A3 | | Biolegend | HeLa | U937 |
| Dy 164 | **CD24** | ML5 | | Biolegend | Kuramochi | U937 |
| Ho 165 | ULBP1 | 170818 | | R&D system | KURAMOCHI | NIH3T3 |
| Er 166 | **CD44** | BJ18 | | Fluidigm | COV362 | OVCAR3 |
| Er 167 | PAX8 | polyclonal | | ProteinTech | Kuramochi | U937 |
| Er 168 | **CD13** | L138 | | BD | U937 | OVCAR4 |
| Tm 169 | **HE4** | EPR4743 | | Abcam | TykNU | U937 |
| Er 170 | Non-p-b Catenin | D13A1 | | CST | Kuramochi | U937 |
| Yb 171 | cPARP | F21-852 | cleaved N214 | BD | U937_Etoposide (20uM, 24hrs) | U937_DMSO (0.05%) |
| Yb 172 | ADAM17 | 111633 | | R&D system | KURAMOCHI | NIH3T3 |
| Yb 173 | **Mesothelin** | MB-G10 | | Rockland | Kuramochi | U937 |
| Yb 174 | ADAM10 | 163003 | | R&D system | KURAMOCHI | NIH3T3 |
| Lu 175 | Total p53 | 1C12 | | CST | Kuramochi | U937 |
| Yb 176 | CD112 | TX31 | | Biolegend | KURAMOCHI | NIH3T3 |

**Table S4**. T**umor and NK receptor ligand antibody panel.** Antibodies against NK receptor ligands (orange highlight) and ADAMs (green highlight). Markers used for X-shift clustering shown in bold.

| Ligand | E vs EV | EV vs V | E vs V | Overall ANOVA |
|---|---|---|---|---|
| ULBP1 | 0.0429 | 0.066 | ns | 0.0052 ** |
| ULBP2 | ns | 0.0001 | ns | 0.0002 *** |
| ULBP3 | ns | 0.0066 | ns | 0.0087 ** |
| ULBP4 | ns | ns | 0.0016 | 0.0023 ** |
| MICA | ns | 0.0001 | 0.0239 | 0.0002 *** |
| ADAM10 | ns | <0.0001 | 0.0066 | <0.0001 **** |
| ADAM17 | ns | <0.0001 | 0.0128 | <0.0001 **** |
| CD111 | ns | 0.0016 | ns | 0.0023 ** |
| CD112 | ns | 0.0016 | 0.0239 | 0.0014 ** |
| CD113 | 0.0429 | 0.0007 | ns | 0.0009 *** |
| CD155 | ns | 0.0033 | ns | 0.0048 ** |
| Nectin4 | ns | <0.0001 | 0.0429 | <0.0001 **** |
| HLA-ABC | ns | ns | ns | ns |
| HLA-E | ns | ns | ns | ns |

**Table S5. Statistical analysis of NK receptor ligand and ADAM expression levels across tumor compartments (For Figure 2D).** p-values shown from one-way ANOVA, paired, ranked with Dunn's multiple comparisons test, adjusted. ns – not significant. p-values: **< 0.01, *** < 0.001, **** < 0.0001.

| Isotope | Antigen | Clone | Vendor | Positive control cell line | Negative control cell line |
|---|---|---|---|---|---|
| Y 89 | CD45 | H130 | Fluidigm | U937 | OVCAR3 |
| In 113 | CD57 | HCD57 | Biolegend | PBMC - CD8+ T cells: CD45RA+CCR7- | PBMC - CD19+ B cells |
| In 115 | HLA-DR | L243 | Biolegend | PBMC - CD14+ Monocytes | PMBC - CD3+ T cells |
| La 139 | 2B4 | C1.7 | Beckman Coulter | NK92 | U937 |
| Pr 141 | CD49d | 9F10 | BD | NKL | NK92 |
| Nd 142 | CXCR3 | G025H7 | Biolegend | HUVEC | PBMC - CD56+ NK cells |
| Nd 143 | VEGF | 23410 | R&D system | NK92_PMA/Iono/Bfa/Mon | NK92_Ethanol (0.2%) |
| Nd 144 | MIP1a/b | 93342 | R&D system | NK92 | NKL |
| Nd 145 | IFNg | B27 | Biolegend | PBMC - CD56+ PMA/Iono/Bfa/Mon | PBMC - CD56+ Ethanol (0.2%) |
| Nd 146 | CD94 | DX22 | Biolegend | PBMC - CD56+ NK cells | PBMC - CD19+ B cells |
| Sm 147 | CD96 | 628211 | R&D system | PBMC - CD56+ NK cells | PBMC - CD19+ B cells |
| Nd 148 | CD34 | 581 | Fluigigm | OVCAR3, K562 | U937 |
| Sm 149 | DNAM1 | TX25 | Biolegend | PBMC - CD56+ NK cells | PBMC - CD19+ B cells |
| Nd 150 | MIP1b | D21-1351 | Fluidigm | PBMC - CD56+ PMA/Iono/Bfa/Mon | PBMC - CD56+ Ethanol (0.2%) |
| Eu 151 | CD107a | H4A3 | Fluidigm | PBMC - CD56+ PMA/Iono/Bfa/Mon | PBMC - CD56+ Ethanol (0.2%) |
| Sm 152 | TNFa | Mab11 | Fluidigm | PBMC - CD56+ PMA/Iono/Bfa/Mon | PBMC - CD56+ Ethanol (0.2%) |
| Eu 153 | EBI3 | B032F6 | Biolegend | NKL_IL12/IL18/Bfa/Mon | NKL_PBS |
| Sm 154 | NKG2D | 1D11 | Biolegend | PBMC - CD56+ NK cells | PBMC - CD19+ B cells |
| Gd 155 | CRACC | 162.1 | Biolegend | PBMC - CD56+ NK cells | PBMC - CD19+ B cells |
| Gd 156 | CD9 | M-L13 | BD | Whole Blood - Platelete CD61+ | Whole Blood - CD3+ T cells |
| Gd 157 | SAP | XLP-1D12 | Cell Signaling | NKL | Ramos |
| Gd 158 | clec12A | 50C1 | Biolegend | PBMC - CD14+ Monocytes | PBMC - CD19+ B cells |
| Tb 159 | CD161 | HP-3G10 | Fluidigm | PBMC - CD56+ NK cells | PBMC - CD19+ B cells |
| Gd 160 | NKp30 | 210845 | R&D system | NK92 | PBMC - CD45+ |
| Dy 161 | Tbet | 4B10 | Fluidigm | PBMC - CD3+ T cells | PBMC - CD19+ B cells |
| Dy 162 | IL-8 | G265-8 | BD | NKL_PMA/Iono/Bfa/Mon (4hrs) | NKL_Ethanol (0.2%) |
| Dy 163 | KIR2DS4 | 179315 | R&D system | PBMC - CD56+ NK cells | PBMC - CD19+ B cells |
| Dy 164 | GMCSF | BVD2-21C11 | Biolegend | NK92_PMA/Iono/Bfa/Mon, 4hrs | NK92_Ethanol (0.2%) |
| Ho 165 | CD16 | 3G8 | Fluidigm | PBMC - CD3- | PBMC - CD4+ T cells |
| Er 166 | TIGIT | MBSA43 | eBioscience | NK92 | NKL |
| Er 167 | IL-10 | JES3-9D7 | Biolegend | NK92_PMA/Iono/Bfa/Mon, 4hrs | NK92_Ethanol (0.2%) |
| Er 168 | KIR2DL1 | 143211 | R&D system | PBMC - CD56+ NK cells | PBMC - CD56- |
| Tm 169 | NKG2A | Z199 | Fluidigm | PBMC - CD56+ NK cells | PBMC - CD56- |
| Er 170 | IL-32 | KU32-52 | Biolegend | PBMC - CD56+ PMA/Iono/Bfa/Mon | PBMC - CD56+ Ethanol (0.2%) |
| Yb 171 | GranzymeB | GB11 | Fluidigm | PBMC - CD56+ NK cells | PBMC - CD19+ B cells |
| Yb 172 | NKp46 | 195314 | R&D system | PBMC - CD56+ NK cells | PBMC - CD19+ B cells |
| Yb 173 | cPARP | F21-852 | BD | U937_Etoposide (20uM, 24hrs) | U937_DMSO (0.05%) |
| Yb 174 | Eomes | WD1928 | eBioscience | PBMC - CD56+ NK cells | PBMC - CD19+ B cells |
| Lu 175 | Perforin | EH12.2H7 | Fluidigm | PBMC - CD56+ NK cells | PBMC - CD19+ B cells |
| Yb 176 | CD56 | NCAM16.2 | Fluidigm | PBMC - CD3-CD14-CD16+ | PBMC - CD4+ T cells |
| Bi 209 | CD47 | CC2C6 | Fluidigm | PBMC - CD45+ | Whole Blood - CD61+ Platelets |

**Table S6. NK cell antibody panel.** Validated antibodies against NK cell receptors and intracellular cytokine expression.

| Cell Line | Catalogue Number | Vendor | Media | Disease Cell Source | Tissue |
|---|---|---|---|---|---|
| OVCAR4 | | Fox Chase Cancer Center | M199; 5% FBS, 1% L-Glu, 1% P/S | High grade ovarian serous adenocarcinoma | Ovary; derived from metastatic site: Ascites |
| Kuramochi | JCRB0098 | JCRB Cell Bank | RPMI; 10% FBS, 1% L-Glu, 1% P/S | Undifferentiated carcinoma | Ascites |
| TYK-nu | JCRB0234.0 | JCRB Cell Bank | EMEM; 10% FBS, 1% L-Glu, 1% P/S | Undifferentiated Carcinoma | Ovary |
| NK-92 | CRL-2407 | ATCC | RPMI, 10% FBS, 1% L-Glu, 1% P/S, 200U/ml IL-2 | 50-year-old Caucasian male with rapidly progressive non-Hodgkin's lymphoma. | Peripheral Blood |
| PC3 | | Brooks Lab, Stanford | FK12; 10%FBS, 1% L-Glu, 1% P/S | Grade IV, adenocarcinoma | Prostate; derived from metastatic site: bone |
| SNU-349 | | Fan Lab, Stanford | RPMI; 10% FBS, 1% L-Glu, 1% P/S | Clear cell renal cell carcinoma | Kidney |
| K562 | CCL-243 | ATCC | IMDM; 10% FBS, 1% P/S | Chronic Myelogenous Leukemia (CML) | Bone Marrow |
| HeLa | CCL-2 | ATCC | DMEM; 10% FBS, 1% L-Glu, 1% P/S | Adenocarcinoma (Cervical) | Cervix |
| A431 | CRL-1555 | ATCC | DMEM; 10% FBS, 1% L-Glu, 1% P/S | Epidermal carcinoma | Skin/Epidermis |
| NCI-H28 | CRL-5820 | ATCC | RPMI; 10% FBS, 1% L-Glu, 1% P/S | Mesothelioma | Lung-pleural effusion |
| HEPG2 | HB-8065 | ATCC | EMEM; 10% FBS, 1% L-Glu, 1% P/S | Hepatocellular Carcinoma | Liver |
| HCT116 | CCL-247 | ATCC | McCoy's; 10% FBS, 1% L-Glu, 1% P/S | Colon Colorectal Carcinoma | Colon |
| MCF7 | HTB-22 | ATCC | EMEM; 0.01 mg/mL human recombinant insulin; 10% FBS, 1% L-Glu, 1% P/S | Adenocarcinoma (Breast) | Mammary Gland |
| MCF10A | CRL-10317 | ATCC | DMEM: F12: 5 % horse serum, 10 µg/mL human insulin, 20ng/mL hEGF, 100ng/mL Cholera toxin, 0.5 µg/mL Hydrocortisone | Fibrocystic disease | Mammary gland, Breast |
| CaCo-2 | HTB-37 | ATCC | DMEM; 20%FBS, 1% L-Glu, 1% P/S | Adenocarcinoma (Colon) | Colon |
| OVCAR3 | HTB-161 | ATCC | McCoy's; 10%FBS, 1% L-Glu, 1% P/S | Adenocarcinoma (Ovarian) | Ovary |
| HCC1937 | CRL-2336 | ATCC | RPMI; 10%FBS, 1% L-Glu, 1% P/S | TNM stage IIB, grade 3, primary ductal carcinoma | Mammary gland; Breast/Duct |
| LoVo | CCL-229 | ATCC | FK12; 10%FBS, 1% L-Glu, 1% P/S | Dukes' Type C, Colorectal Adenocarcinoma | Colon |
| A549 | CCL-185 | ATCC | FK12; 10%FBS, 1% L-Glu, 1% P/S | Carcinoma | Lung |
| OVSAHO | JCRB1046 | JCRB Cell Bank | RPMI; 10%FBS, 1% L-Glu, 1% P/S | Ovarian Carcinoma | Abdominal Metastasis |
| OVKATE | JCRB1044 | JCRB Cell Bank | RPMI; 10%FBS, 1% L-Glu, 1% P/S | Ovarian Carcinoma | Ovary |
| SNU-119 | | Seoul National University - Korea | RPMI; 10%FBS, 1% L-Glu, 1% P/S | Cystadenocarcinoma, serous, papillary | Ovary; Ascites |
| JHOS-2 | RBRC-RCB1521 | RIKEN BRC Cell Bank | DMEM:Ham F12:10% FBS,0.1nM NEAA, 1% P/S | Serous adenocarcinoma of ovary | Ovary |
| JHOM-2B | RBRC-RCB1682 | RIKEN BRC Cell Bank | DMEM:HamF12:10% FBS,0.1nM NEAA, 1% P/S | Mucinous tubular adenocarcinoma | Ovary |
| COV362 | 7071910 | Public Health England | DMEM; 10% FBS, 1% L-Glu, 1% P/S | Endometroid Carcinoma | Ovary |
| DLD-1 | HD PAR-008 | Horizon Discovery | RPMI; 10%FBS, 1% L-Glu, 1% P/S, 25mM sodium bicarbonate, | Colorectal adenocarcinoma Dukes Type C | Colon |
| COV318 | 7071903 | Sigma Aldrich | DMEM; 10% FBS, 1% L-Glu, 1% P/S | Ovarian epithelial-serous carcinoma | Ascites |

**Table S7. HGSC and non-HGSC tumor cell lines used for this study.** All cell lines were authenticated by STR sequencing. FBS- fetal bovine serum, P/S – penicillin/streptomycin L-Glu – L-glutamine, NEAA – non-essential amino acids.
